# *Helicobacter pylori* biofilm cells are metabolically distinct, express flagella, and antibiotic tolerant

**DOI:** 10.1101/728766

**Authors:** Skander Hathroubi, Julia Zerebinski, Karen M. Ottemann

## Abstract

Biofilm growth protects bacteria against harsh environments, antimicrobials, and immune responses. *Helicobacter pylori* is a bacterium that has a robust ability to maintain colonization in a challenging environment. Over the last decade, *H. pylori* biofilm formation has begun to be characterized, however, there are still gaps in our understanding about how this growth mode is defined and its impact on *H. pylori* physiology. To provide insights into *H. pylori* biofilm growth properties, we characterized the antibiotic susceptibility, gene expression, and genes required for biofilm formation of a strong biofilm-producing *H. pylori*. *H. pylori* biofilms developed complex 3D structures and were recalcitrant to multiple antibiotics. Disruption of the protein-based matrix decreased this antibiotic tolerance. Using both transcriptomic and genomic approaches, we discovered that biofilm cells demonstrated lower transcripts for TCA cycle enzymes but higher ones for hydrogenase and acetone metabolism. Interestingly, several genes encoding for the natural competence Type IV secretion system 4 (*tfs4*) were up-regulated during biofilm formation along with several genes encoding for restriction-modification (R-M) systems, suggesting DNA exchange activities in this mode of growth. Flagella genes were also discovered through both approaches, consistent with previous reports about the importance of these filaments in *H. pylori* biofilm. Together, these data suggest that *H. pylori* is capable of adjusting its phenotype when grown as biofilm, changing its metabolism and elevating specific surface proteins including those encoding *tfs4* and flagella.

## INTRODUCTION

More than half of the world’s population is infected with *Helicobacter pylori*, which makes it one of the most common bacterial infections affecting humans (Testerman and Morris 2014). *H. pylori* is the main cause of chronic gastritis, duodenal and gastric ulcers, and significantly increases the risk of developing gastric cancer and mucosal-associated-lymphoid type (MALT) lymphoma. Gastric cancer kills over 700,000 people per year, with an estimated cost associated with peptic ulcer disease of 6 billion dollars in the US alone (Parkin, Bray et al. 2005, Testerman and Morris 2014).

Treating *H. pylori* infection remains challenging (Jones, Cha et al. 2008, Vakil and Vaira 2013), and when eradication strategies fail, *H. pylori* can persist for the lifetime of the host. Several mechanisms are involved in the persistence and antibiotic tolerance of *H. pylori* (Jones, Cha et al. 2008), however one of them, biofilm formation, remains understudied. Biofilms are considered as the natural mode of bacterial growth (Hall-Stoodley, Costerton et al. 2004). Within a biofilm, cells are often highly resistant to environmental stress, as well as recalcitrant to the action of antibiotics and the host immune system. These properties make biofilm growth one of the major global challenges to modern medicine (Costerton, Stewart et al. 1999, Hathroubi, Mekni et al. 2017). Several investigations have shown that *H. pylori* cells can grow as biofilm *in vitro* (Stark, Gerwig et al. 1999, Yonezawa, Osaki et al. 2010, Yonezawa, Osaki et al. 2011, Servetas, Carpenter et al. 2016, De la Cruz, Ares et al. 2017, Hathroubi, Zerebinski et al. 2018) as well as *in vivo* (Carron, Tran et al. 2006, Coticchia, Sugawa et al. 2006, Cammarota, Branca et al. 2010) and make multicellular complex surface-attached communities. These studies suggest that biofilm formation may be a significant contributor to *H. pylori* persistence.

Although the molecular mechanisms that drive *H. pylori* biofilm formation are not fully elucidated, several factors associated with *H. pylori* biofilm have been identified using high throughput strategies such as genomics, transcriptomics, and proteomics (Shao, Sun et al. 2013, Wong, Ng et al. 2016, Hathroubi, Servetas et al. 2018, Hathroubi, Zerebinski et al. 2018). Genes expressed to high levels in the biofilm include several genes or products of the cytotoxin-associated gene pathogenicity island (*cag*PAI) which encode for a type IV secretion system, the *cag*PAI substrate CagA, as well as diverse membrane proteins (Shao, Sun et al. 2013, Wong, Ng et al. 2016). A recent RNA sequencing comparison between planktonic and biofilm cells of *H. pylori* strain SS1 found that planktonic and biofilm cell populations displayed distinct transcriptomic profiles, with genes related to cell envelope and the flagellar apparatus up-regulated in biofilm cells compared to their planktonic counterparts (Hathroubi, Zerebinski et al. 2018). Surprisingly, flagellar filaments were observed in high abundance in the biofilm and shown to promote biofilm formation. Biofilm cells were also less metabolically active than planktonic cells and several genes associated with stress responses were up-regulated in biofilm cells, such as those encoding for peptidoglycan deacetylase (*pgdA*), DNA recombination (*recR*), and heat shock transcriptional regulators HspR and HrcA (Hathroubi, Zerebinski et al. 2018). *H. pylori* SS1 required low nutrient conditions to produce a pronounced biofilm (Hathroubi, Zerebinski et al. 2018). In contrast, *H. pylori* strain G27 readily forms biofilms under nutrient poor and nutrient rich conditions such as those used routinely in the laboratory (Hathroubi, Zerebinski et al. 2018, Windham, Servetas et al. 2018). *H. pylori* G27 therefore appears to be a strong biofilm-producing strain, consistent with its previous use as a model for studying *H. pylori* biofilm formation (Anderson, Huang et al. 2015, Gaddy, Radin et al. 2015, Servetas, Carpenter et al. 2016, Servetas, Doster et al. 2018, Windham, Servetas et al. 2018). Because the pathways for biofilm formation had not yet been characterized in *H. pylori* strain G27, we performed transcriptomic and genomic approaches to unravel genes associated with the biofilm mode of growth in this strong biofilm-forming *H. pylori*, and characterized how biofilm growth affected antibiotic susceptibility.

## METHODS

### Bacterial strain and growth conditions

This study employed *H. pylori* wild type and clinical strains, as well as several mutants generated using a transposon library that was constructed in strain G27 and generously provided by Nina Salama (Baldwin, Shepherd et al. 2007) (Table 1). Strains were grown on Colombia Horse Blood Agar (CHBA) (Difco), containing: 0.2%*β*-cyclodextrin, 10 μg of vancomycin per ml, 5 μg of cefsulodin per ml, 2.5 U of polymyxin B per ml, 5 μg of trimethoprim per ml, and 8 μg of amphotericin B per ml (all chemicals are from Thermo Fisher or Gold Biotech), or Brucella broth (Difco) containing 10% fetal bovine serum (BB10; Gibco/BRL). Cultures were grown under microaerobic conditions (10% CO_2_, 5% O_2_, 85% N_2_) at 37°C. For antibiotic resistance marker selection, bacterial media were additionally supplemented with 25μg of chloramphenicol (Cm) per ml or 75μg kanamycin (Km) per ml.

**Table 1.** Strains used for this in the present study.

| *Strain* | **KO reference no.** | **Genotype or description** | **Reference and/or source(s)** |
| --- | --- | --- | --- |
| ***G27*** | KO379 | WT strain | N. Salama |
| ***SS1*** | KO457 | WT strain | J. O'Rourke |
| ***J99*** | KO479 | WT strain | N. Salama |
| ***26695*** | KO668 | WT strain | ATCC 700392 |
| ***X472AL*** | KO612 | WT strain | D. Berg |
| ***M6*** | KO613 | WT strain | E. Joyce, A. Wright |
| ***CYP34 (CYP3401)*** | KO637 | WT strain | D. Berg |
| ***HP43504*** | KO669 | WT/clinical isolate | NTCT 11637 |
| ***HP 100-1*** | KO763 | WT/clinical isolate | J. Solnick |
| ***HP 12-1*** | KO760 | WT/clinical isolate | J. Solnick |
| ***HP 28-1*** | KO761 | WT/clinical isolate | J. Solnick |
| ***HP116-1*** | KO764 | WT/clinical isolate | J. Solnick |
| ***HP 125-2*** | KO765 | WT/clinical isolate | J. Solnick |
| ***HP 208-2*** | KO767 | WT/clinical isolate | J. Solnick |
| ***G27 ΔHPG27_166<sup>#</sup>*** | KO1635 | *ΔHPG27_166:aphA3* | This study |
| ***G27 ΔHPG27_526<sup>#</sup>*** | KO1637 | *ΔHPG27_526::cat* | This study |
| ***G27 ΔhydE<sup>#</sup>*** | KO1638 | *ΔhydE::cat* | This study |
| ***G27 ΔHPG27_1233<sup>#</sup>*** | KO1639 | *ΔHPG27_1233::cat* | This study |
| ***G27 ΔHPG27_1066<sup>#</sup>*** | KO1640 | *ΔHPG27_1066::cat* | This study |
| ***G27 ΔacsA<sup>#\*</sup>*** | KO1641 | *ΔacsA::cat* | This study |
| ***G27 ΔacxAB<sup>#</sup>*** | KO1642 | *ΔacxAB::cat* | This study |
| ***G27 ΔHPG27_715<sup>#</sup>*** | KO1644 | *ΔHPG27_715::cat* | This study |

<sup>#</sup>All transformations were performed using natural transformation using plasmid constructs (pTwist\_Amp) in which the gene's coding sequence was completely or partially removed and replaced by a kanamycin or chloramphenicol-resistance cassette using allelic exchange.
\*Mutant *ΔacsA* showed a slight defect in growth.
WT, wild-type; *aphA3*, kanamycin resistant; *cat*, chloramphenicol resistant; ATCC, American Type Culture Collection.

### Biofilm formation

Biofilm formation assays were carried as described previously, with slight modifications (Hathroubi, Zerebinski et al. 2018). *H. pylori* strains were grown overnight with shaking in BB10, diluted to an OD600 of 0.15 with fresh BB10 media and then used to fill triplicate wells of a sterile 96-well polystyrene microtiter plate (Costar, 3596). Following static incubation of three days under microaerobic conditions, culture medium was removed by aspiration and the plate was washed twice using 1x Phosphate-buffered saline (PBS). The wells were then filled with 200μL of crystal violet (0.1 %, wt/vol), and the plate was incubated for 2 min at room temperature.

After removal of the crystal violet solution by aspiration, the plate was washed twice with PBS and air dried for 20 min at room temperature. To visualize biofilms, 200 μL of ethanol (70%, vol/vol) was added to the wells and the absorbance at 590 nm was measured.

### Antimicrobial susceptibilities

The antimicrobial susceptibilities for amoxicillin, clarithromycin and tetracycline were determined using E-test strips for each antibiotic (BioMérieux). Overnight cultures of *H. pylori* were adjusted to an optical density at 600nm of 0.5 and cultures were inoculated on CHBA plate. E-test strips were aseptically placed onto the plates. After 72 h of incubation, MIC was recorded. The E-test minimal inhibitory concentration (MIC) was interpreted as the point at which the growth intersected the strip. The experiment was conducted in triplicate for each antibiotic.

### Planktonic and Biofilm antimicrobial susceptibility

Biofilms were cultured as described above in 96-well plates. After 2 days of incubation, the supernatant was removed by aspiration and 200 μL of media containing the desired antibiotic concentration was added to each well. Wells containing only the bacterial inoculum were used as positive controls and wells containing only media with the antibiotic was used as negative control. The plates were incubated for 24 hours and then the biofilms were washed, stained and quantified as described above. Alternatively, biofilms exposed or not to clarithromycin were washed and resuspended in PBS and subjected to colony forming unit (CFU) determination. For planktonic cultures, overnight, 2-days or 3-days *H. pylori* cultures where exposed to the desired antibiotic concentration for 24 hours in both static and shaking conditions and then plated for enumeration by CFU counting.

### Biofilm dispersion assay

Enzymatic dispersion of established biofilm was performed as described previously (Hathroubi, Zerebinski et al. 2018, Windham, Servetas et al. 2018). Briefly, biofilms were grown as above, and after 3-days of incubation, supernatants were replaced with either fresh media, fresh media containing proteinase K (200ug/mL to 0.05ug/mL, Sigma-Aldrich), or fresh media containing proteinase K (200ug/mL to 0.05 ug/mL) and clarithromycin (MIC or 128x MIC). For control wells, supernatants were replaced with fresh media. Wells were treated for 6 hours or 24 hours under the desired conditions, and biofilms were washed, stained and quantified as described above.

### Biofilm and Planktonic growth conditions for transcriptomic analysis

Planktonic cells were grown in BB10 as above with constant shaking for 24h at 37°C under microaerobic conditions. After incubation, planktonic cells were harvested by centrifugation (10min, 4900 rpm) and resuspended in 1mL of Trizol Max from the Ambion Bacterial Enhancement Kit (Ambion, Life technology, Carlsbad, CA, USA). For biofilm cells, *H. pylori* G27 was grown in 6-well plates (Costar) in BB10 for three days, the time required for *H. pylori* to develop a notable biofilm. After incubation, cells grown in biofilm condition, were scrapped of from the surface with a sterile cell scraper, harvested by centrifugation (10min, 4900 rpm), washed twice with PBS, and resuspended in 1mL of Trizol Max.

### RNA extraction and library construction

Total RNA from *H. pylori* was extracted using the Trizol Max Bacterial Enhancement Kit (Ambion, Life Technology, Carlsbad, CA, USA) as described by the manufacturer. RNA was further purified and concentrated using an RNAeasy Kit (Qiagen). rRNA was removed using RiboZero magnetic kit (Illumina). Sequencing libraries were generated using NEBNext Ultra^TM^ Directional RNA library Prep Kit for Illumina (NEB, USA). cDNA library quality and amount were verified using Agilent Bioanalyzer 2100 system (Agilent technologies, CA, USA) and then sequenced using Illumina NextSeq Mid-Output (UC Davis Genome Center).

### Transcriptomic analysis

RNA-seq data were analyzed using CLC Genomics Workbench (version 11.0, CLC Bio, Boston, MA, USA). All sequences were trimmed, and forward and reverse sequenced reads generated for each growth state (biofilm vs planktonic; three biological replicates for each condition) were mapped against the G27 reference genome (Baltrus, Amieva et al. 2009) to quantify gene expression levels for each experimental condition. The expression value was measured in Reads per Kilobase Per Million Mapped Reads (RPKM). Genes were considered as differentially expressed when log2 (fold change) was above 1 or below −1 and with statistical significance (*P*-value <0.05, false discovery rate (FDR < 0.001).

### Quantitative PCR

In order to validate RNA-seq data, qRT-PCR was performed to quantify the transcription of six selected genes (two up-regulated; *virC1* and *virB11*, two down-regulated; *fdxB* and HPG27_526 and two non-differentially expressed genes; *lctP* and *hopC*) Total RNA from 3 independent experiments was obtained and used for qRT-PCR. Primers were designed using Primer-Blast (https://www.ncbi.nlm.nih.gov/tools/primer-blast/) and are listed in **Table 5**. The same amount of total RNA (1μg/μL) was reverse transcribed using the LunaScript^TM^ RT SuperMix Kit (NEB) and qPCR reactions were prepared using Luna Universal qPCR Master Mix kit (NEB) The run was performed in Connect^TM^ Thermal Cycler (Bio-Rad) with the following cycling parameters: 2 min at 95°C, followed by 39 cycle of 30 s at 95°C, 30 s at 60°C, and 30 s at 72°C. Relative expression of each gene was normalized to that of the 16S gene, whose expression was consistent throughout the different conditions. Quantitative measures were made using the 2^−^*^ΔΔ^*^C^T method (Rao, Huang et al. 2013). Three biological replicates and two technical replicates of each condition were performed.

### Screening approach for biofilm-defective mutants

A *Tn*-7 based transposon-based mutant library previously generated in *H. pylori* G27 (Baldwin, Shepherd et al. 2007) was grown overnight in BB10 supplemented with 25 μg of chloramphenicol, diluted to an OD600 of 0.15 and then used to fill duplicate wells of a 6-well plate (Costar® 3516, Corning, Corning, NY, USA). Media containing non-attached, planktonic bacteria and/or potential biofilm defective mutants was removed after different incubation lengths (i.e. 1, 2, 3, 24, 48 or 72 hours), and transferred to a new sterile 6-well plate. The procedure was repeated 2 times. After 72 hours, this enriched biofilm-defective sample was plated on CHBA media and individual colonies were isolated and stored at −80°C. A total of 97 potential biofilm defective mutants was isolated from this library, and subsequently verified for abnormal biofilm formation using the biofilm assay described above.

### Identification of transposon interrupted genes

Nested PCR was conducted for the identification of the disrupted gene in the potential defective-biofilm mutant obtained from the screening. As described before (Salama, Shepherd et al. 2004), for the first round of PCR we used random primers with a constant 5’ tail regions in tandem with a transposon specific primer (Upstream or Downstream) (Supplemental Table 2). For the second round, a primer specific to the transposon was utilized with a primer complementary to the tail region of the original random primer. Therefore, the final products contain a portion of the transposon along with surrounding genomic information. These PCR products were sequenced (Sequetech, Mountain View, CA) and then BLASTed against the G27 genomic sequence at the UCSC Microbial Genome Browser (www.http://microbes.ucsc.edu).

### Design and construction of mutants

Clean gene knockout constructs were synthesized by Twist Bioscience (San Francisco, CA). For each target, the gene’s coding sequence was fully replaced by allelic exchange with a chloramphenicol resistance cassette (*cat*) or a kanamycin resistance cassette (*aphA3*). All constructs were cloned in the pTwist Amp high copy vector and transformed into *Escherichia coli* DH10B. Purified vector containing the construct (2 to 10 ug) were used to transform *H. pylori* G27 WT. Mutants were selected on chloramphenicol or kanamycin-containing plates, and colony purified.

### Confocal laser scanning microscopy

Biofilms of *H. pylori* G27 and different defective-biofilm mutants were prepared as described above using BB10 media, however, for confocal laser scanning microscopy (CLSM), µ-Slide 8-well glass bottom chamber slides (ibidi, Germany) were used instead of 96-well microtiter plates. Three-day-old biofilms were stained with FilmTracer*™* FM*^®^*1–43 (Invitrogen), BOBO-3 (Invitrogen), FilmTracer*™* SYPRO*®* Ruby biofilm matrix stain (Invitrogen), or FilmTracer LIVE/DEAD biofilm viability kit (Invitrogen) according to the manufacturer’s instructions. Stained biofilms were visualized by CLSM with an LSM 5 Pascal laser-scanning microscope (Zeiss) and images were acquired using Imaris software (Bitplane). Biomass analysis of biofilm was carried out using FM*^®^*1–43 stained z-stack images (0.15μm thickness) obtained by CLSM from randomly selected areas. The biomass of biofilms was determined using COMSTAT (Heydorn, Nielsen et al. 2000).

### Scanning electron microscopy

*H. pylori* wild-type G27 and defective-biofilm mutants were grown on glass coverslips in 6-well plates (Costar) by dispersing 4 mL of a culture diluted to OD 0.15 in BB10 into the coverslip containing wells. The plate was incubated for 3-days under microaerobic conditions. Biofilms formed on the surface of the coverslips were washed twice with PBS and fixed with 2.5% glutaraldehyde for 1h at room temperature. Samples were then dehydrated with ethanol, critically point dried, sputtered with ∼20 nm of gold and imaged in a FEI Quanta 3D Dual beam SEM operating at 5 kV and 6.7 pA.

### Statistical analysis

Collected biofilm data were analyzed statistically using GraphPad Prism software (version 7, GraphPad Software Inc., San Diego, CA) by application of Wilcoxon-Mann-Whitney test or one-way ANOVA with Bartlett’s test. *p* < 0.05 or < 0.01 were considered reflecting statistically significant differences.

## RESULTS

### Biofilm is a common trait among H. pylori and strain G27 is a strong biofilm-forming strain

Several studies have shown multiple *H. pylori* strains form biofilms, including strain G27 (Cole, Harwood et al. 2004, Williams, McInnis et al. 2008, Yonezawa, Osaki et al. 2010, Bessa, Grande et al. 2013, Anderson, Huang et al. 2015, Servetas, Doster et al. 2018, Windham, Servetas et al. 2018). However, these multiple strains had not been compared side-by-side. Therefore, in this study, we first compared the biofilm-forming ability of strain G27 with several *H. pylori* reference and clinical strains using the microtiter plate biofilm assay, growth in BB10, and crystal violet staining. All strains were able to adhere to the plastic surfaces and grow as multicellular aggregates, suggesting that biofilm formation is a common *H. pylori* trait (Fig. 1). This assay revealed significant variations in biofilm biomass between strains. Some strains, such as G27 and X47, were “high” biofilm producers while others, including the common mouse-infecting strain SS1 and some clinical isolates (i.e. H100-1, H12-1 and H125-2), were “low” biofilm producers (Fig. 1). While the reason for this variation is not yet clear, we decided to focus on the properties of the high biofilm producer G27.

**Figure 1.**
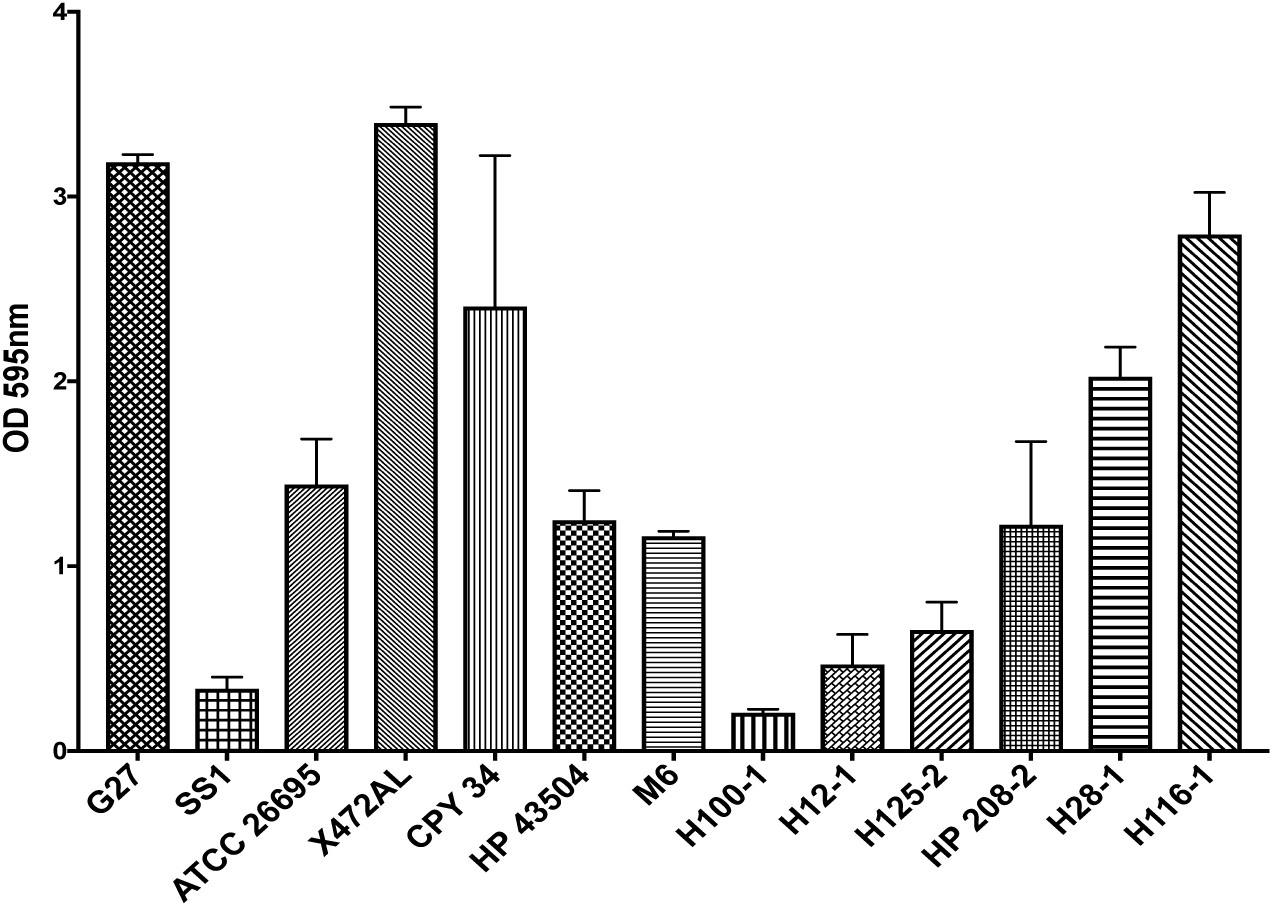
*H. pylori* G27 is a high biofilm forming strain. Biofilm formation of *H. pylori* strains were assessed using the microtiter plate biofilm assay, with strains grown for three days in BB10. Results represent the crystal violet absorbance at 595nm, which reflects the biofilm biomass. Experiments were performed three independent times with at least 6 technical replicates for each. Error bars represent standard error of the mean

It is worth noting that strain G27 produced varied forms of biofilm in addition to the biofilm formed on plastic surface at the bottom of wells. Specifically, we noted a ring at the air-liquid interface and a thin, extremely fragile pellicle floating at the air-liquid interface, as seen by others (Servetas, Carpenter et al. 2016). This pellicle was not observed in other strains including SS1 (Hathroubi, Zerebinski et al. 2018). This observation suggests that some *H. pylori* might be able to form multiple types of biofilm aggregates, however for this work we focused on those formed at the bottom surface of the wells.

### G27 biofilm consists of aggregation of mainly live cells and interacted to each other by flagella filaments and pili-like structures

To visualize *H. pylori* grown as surface-attached biofilms in more detail, confocal scanning laser microscopy (CSLM) and scanning electron microscopy (SEM) were performed. Live-dead staining revealed that *H. pylori* G27 3-day biofilm consists of flat, uniform layers of bacteria with a thickness of 15-20 μm (Fig. 2A). Live populations (green) were predominant throughout the biofilm compared to dead population (red) with a live/dead ratio estimated at 1.17±14 (Fig. 2A).

**Figure 2.**
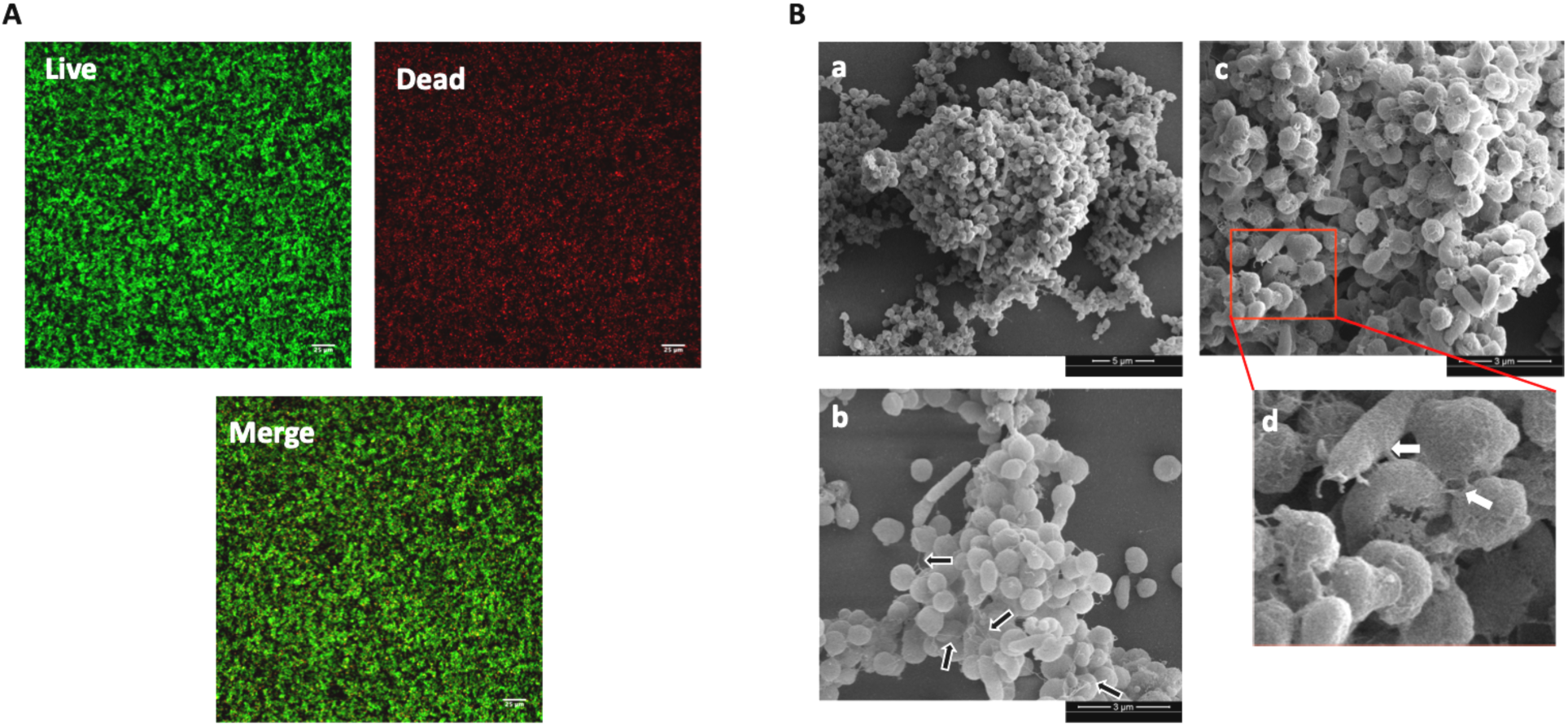
*H. pylori* G27 Biofilm characterization. **(A)** Confocal scanning laser microscopy (CSLM) images (top view) of 3-day old *H. pylori* G27 biofilms stained with LIVE/DEAD stain, resulting in live and dead bacteria appearing as green or red, respectively**. (B)** SEM images show cell-to-cell interactions through flagella-like filaments and pili-like structures within a 3-day old *H. pylori* G27 biofilm. Biofilms were grown in BB10 media. High magnification in (c) is derived from the red-boxed area of (b). Black arrows indicate flagella filaments and white arrows indicate the presence of pili-like structures connecting cells together.

We next applied scanning electron microscopy (SEM) to gain more insight into the biofilm structure. SEM images showed 3D complex structures composed of aggregated bacteria attached to the surface and together. These aggregates contained mainly coccoid forms along with some bacilli (Fig. 2B), as reported previously for *H. pylori* strain SS1 (Hathroubi, Zerebinski et al. 2018). Rod-shaped cells were mostly characterized by straight rod morphology of about 2-3 μm long by 0.5 μm wide, whereas cocci had a diameter around 0.4-0.6 μm (Fig. 2B). Biofilms contained many filaments including ones that look like the flagellar filaments described previously as structurally important in *H. pylori* biofilm (Hathroubi, Zerebinski et al. 2018). Interestingly, pili-like structures were also detected in biofilms at cell-to-cell contacts, however their exact nature and role remain to be determined (Fig. 2B).

### G27 biofilm formation is time and pH dependent but not affected by FBS variation

*H. pylori* G27 formed biofilms in BB media supplemented with either 10% or 2% FBS (Fig. 1 and data not shown), consistent with previous reports that the G27 biofilm is relatively stable and unaffected by slight variations of growth media (Windham, Servetas et al. 2018). By monitoring the strain G27 biofilm over several days, we concluded that 24-hours of growth was required to develop a noticeable biofilm and 72-hours of growth was needed to reach the peak of biomass (Fig. S1A). Biofilm formation was influenced by some environmental parameters including pH. We observed that biofilm formation gradually decreased as the pH was lowered from neutral to pH 3 (Fig. S1B). However, once formed, biofilms were not affected by exposure to moderately low pH (5.5) but were eliminated by exposure to pH3 (Data not shown). These results suggest that pH might be a key determinant of biofilm formation. For our subsequent work, we used biofilm growth in BB media supplemented with 10% FBS, pH 7.0, and 2 or 3 days of incubation as our standard conditions.

### Biofilm cells demonstrate high levels of antibiotic tolerance compared to planktonic cells

It is well known that bacteria within biofilm can be 10 to 10,000 times more tolerant to antibiotics than their planktonic counterparts (Hoiby, Bjarnsholt et al. 2010, Stewart 2015, Hathroubi, Mekni et al. 2017). However, the impact of biofilm mode of growth on susceptibility to antimicrobial agents in *H. pylori* has not been well documented. Previous studies have shown that *H. pylori* clinical strains TK1402 and TK1049 increased levels of tolerance and resistance to clarithromycin of up to 8 to 16-fold times the minimum inhibitory concentration (MIC) as compared to planktonic grown cells (Yonezawa, Osaki et al. 2013, Yonezawa, Osaki et al. 2019). We thus decided to investigate this phenomenon in the strong-biofilm forming strain G27 and determine biofilm and planktonic cells susceptibility to three clinically relevant antibiotics: the macrolide clarithromycin, the *β*-lactam amoxicillin, and tetracycline. Forty-eight-hour old *H. pylori* G27 biofilms were treated with either amoxicillin, tetracycline, or clarithromycin for 24-hours and biofilm biomass was then evaluated using the crystal violet assay. These assays revealed that pre-formed biofilms were highly tolerant to all antibiotics at up to 128-times the established MIC (Fig. 3). Biofilms exposed to amoxicillin and tetracycline remained practically unchanged even when exposed to 128-times the MIC (Fig. 3). Exposure to clarithromycin did significantly decrease the biofilm biomass compared to untreated controls, however, the biofilms were not eliminated and remained detectable at concentrations reaching up to 128 times the MIC (Fig. 3).

**Figure 3.**
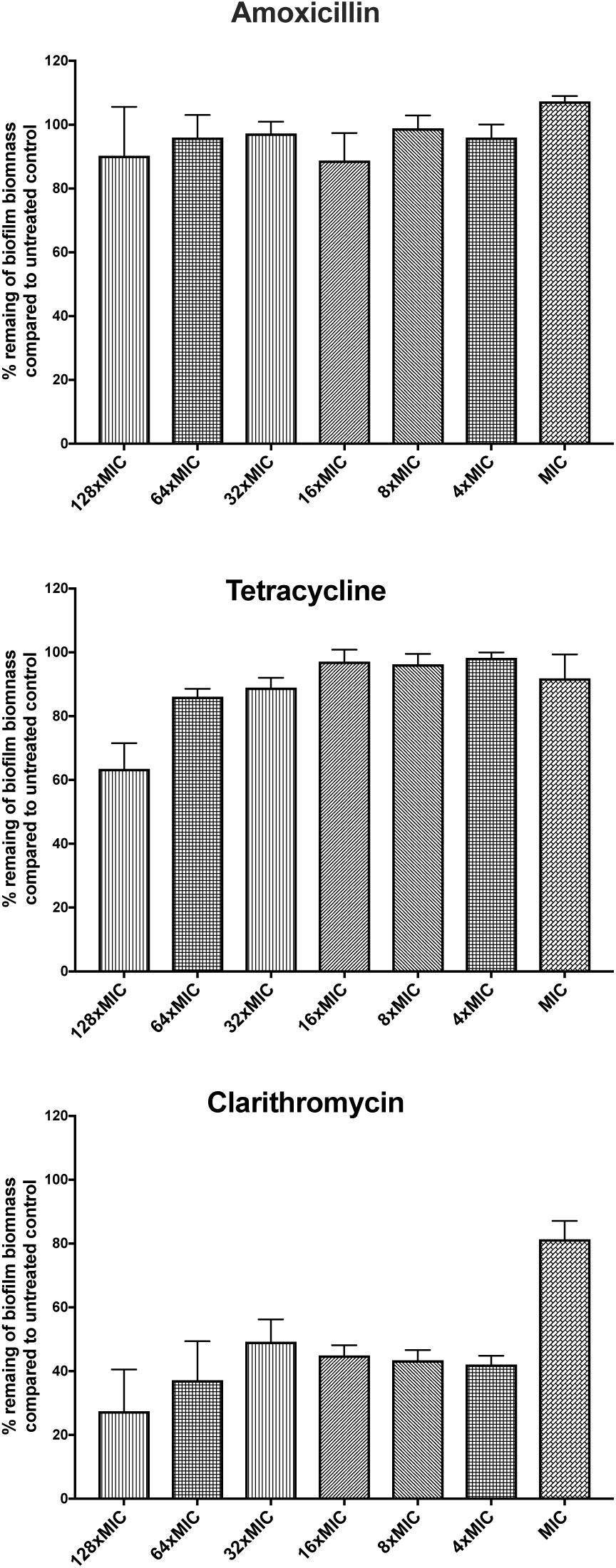
*H. pylori* biofilm growth is associated with high level tolerance towards amoxicillin, tetracycline and clarithromycin. The percentage of remaining *H. pylori* G27 biofilm biomass after exposure to different concentration of amoxicillin, tetracycline or clarithromycin compared to untreated control. Control biofilms were exposed to fresh media without antibiotic. Biofilm formation of *H. pylori* strain were assessed using the microtiter plate biofilm assay at different time points. Experiments were performed three independent times with at least 8 technical replicates for each. Error bars represent standard error of the mean.

The viability of planktonic and biofilm cells after exposure to either clarithromycin, amoxicillin or tetracycline was determined by plating the samples to analyze live cells by colony forming units (CFU) (Fig. 4A). As expected, cells grown in planktonic conditions for 48 hours were completely killed by exposure to either 1-times or 128-times MIC levels of each antibiotic, resulting in no viable cells (Fig. 4A). Conversely, cells grown as biofilms tolerated exposure to amoxicillin or tetracycline up to 128-times MIC, with only 12.5% and 7.5% of biofilm cell population killed (Fig. 4A). Clarithromycin was more able to kill biofilm cells with exposure at 128-times MIC eliminating approximately half the population.

**Figure 4.**
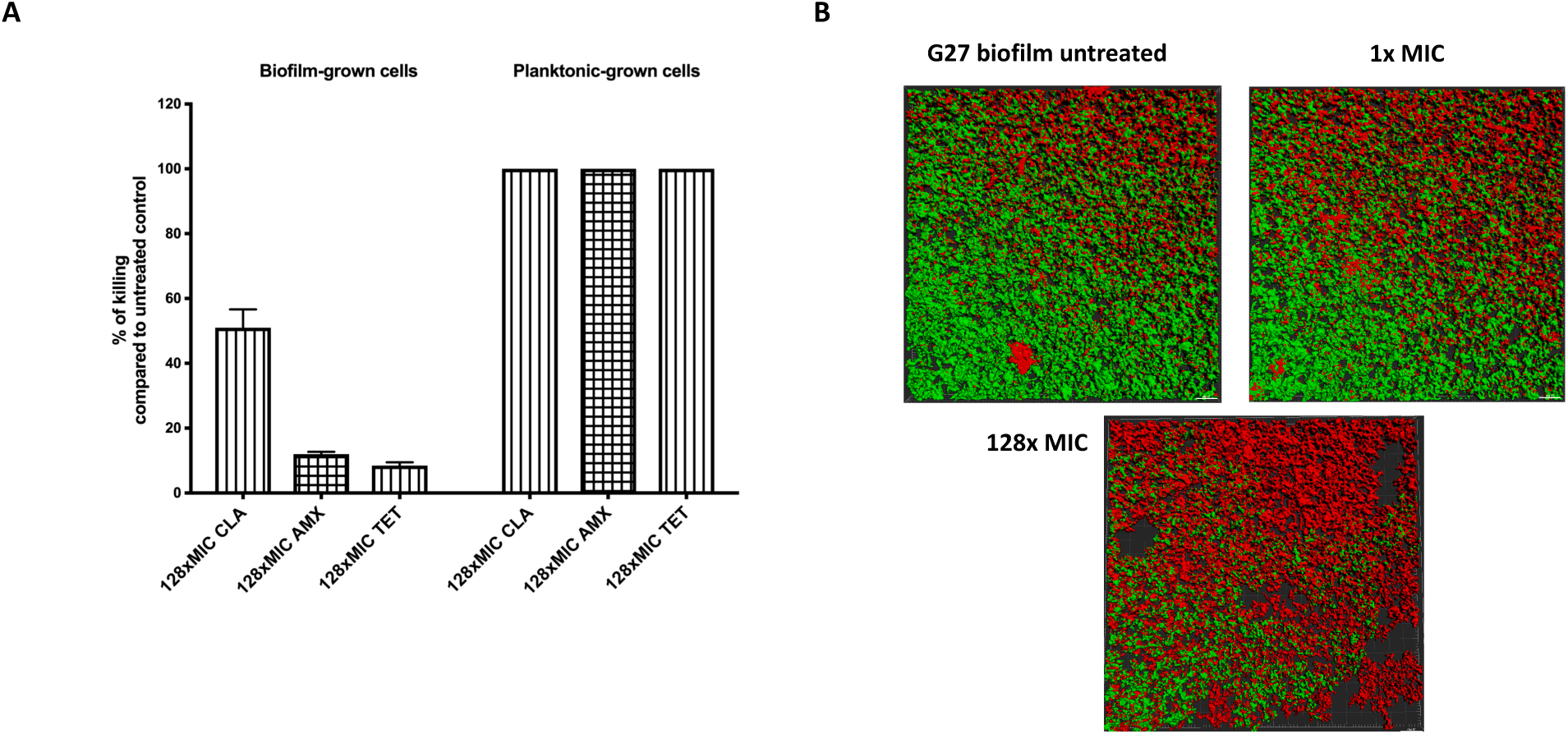
Viability of *H. pylori* cells grown as biofilm or planktonic after exposure to antibiotics. **(A)** *H. pylori* G27 grown in biofilm (48 hours) or planktonic (48 hours) conditions and then treated or not with either clarithromycin (CLR), amoxicillin (AMX) and tetracycline (TET) for an additional 24 hours. Data represent percentage of killing (Colony forming units (CFU) per mL) compared to untreated control. **(B)** Live and dead biomasses in *H. pylori* G27 biofilm treated or not with clarithromycin. Biofilms were visualized by CLSM with LIVE/DEAD viability strain. Viable cells exhibit green fluorescent, whereas dead and damaged cells exhibit red fluorescence. These images are three-dimensional (3D) reconstructions from an average of 12 image stacks using Imaris software.

We further evaluated the impact of clarithromycin on pre-formed biofilms using confocal microscopy (Fig. 4B). As shown above, biofilms contain both live and dead cells (Fig.2A). However, despite a treatment with up to 128-times of the MIC of clarithromycin, live populations were still observed. (Fig. 4B). This result is similar to that obtained with CFU analysis, in that clarithromycin killed some cells but not all. Together, these data suggest that when strain G27 grows as a biofilm, it becomes highly tolerant to antimicrobials.

### Biofilm matrix-associated protein restrict clarithromycin penetration

Biofilms can constitute a physical and/or chemical diffusion barriers to antimicrobial penetration (Stewart 2015, Hathroubi, Mekni et al. 2017). *H. pylori* is known to use a matrix composed mainly of eDNA and proteins (Grande, Di Giulio et al. 2011, Grande, Di Marcantonio et al. 2015, Hathroubi, Zerebinski et al. 2018, Windham, Servetas et al. 2018). We thus tested whether the extracellular matrix of *H. pylori* can limit the penetration of antibiotics, focusing on the most commonly used *H. pylori* antibiotic, clarithromycin. To this end, we used proteinase K, which was previously shown to successfully disperse pre-formed *H. pylori* G27 biofilms (Windham, Servetas et al. 2018). We first determined doses of proteinase K that did not entirely eradicate pre-formed biofilms, so that biofilms structure would be only partially destabilized but possibly allow diffusion of clarithromycin (Fig. 5). Doses between 5 and 0.5 μg/mL of proteinase K did not disperse the biofilm (Fig. 5A) and did not affect the viability of strain G27 (data not shown). Combining proteinase K (0.5 μg/mL) with clarithromycin at 128-times the MIC demonstrated a significant higher biofilm inhibition compared to a treatment with clarithromycin alone (Fig. 5B). This outcome suggests that some target of proteinase K, such as the extracellular matrix or outer-membrane proteins, limits the exposure to clarithromycin. We did note that these combined treatments did not completely eradicate *H. pylori* biofilms, so it’s possible that other mechanisms associated with biofilm growth in *H. pylori* such as low metabolism (Hathroubi, Zerebinski et al. 2018) and overexpression of efflux pumps (Yonezawa, Osaki et al. 2013) could be implicated in this phenomenon. Further experiments will be needed to elucidate mechanisms behind antimicrobial tolerance in *H. pylori* biofilms, but our data suggested that protein components play some part in this process.

**Figure 5.**
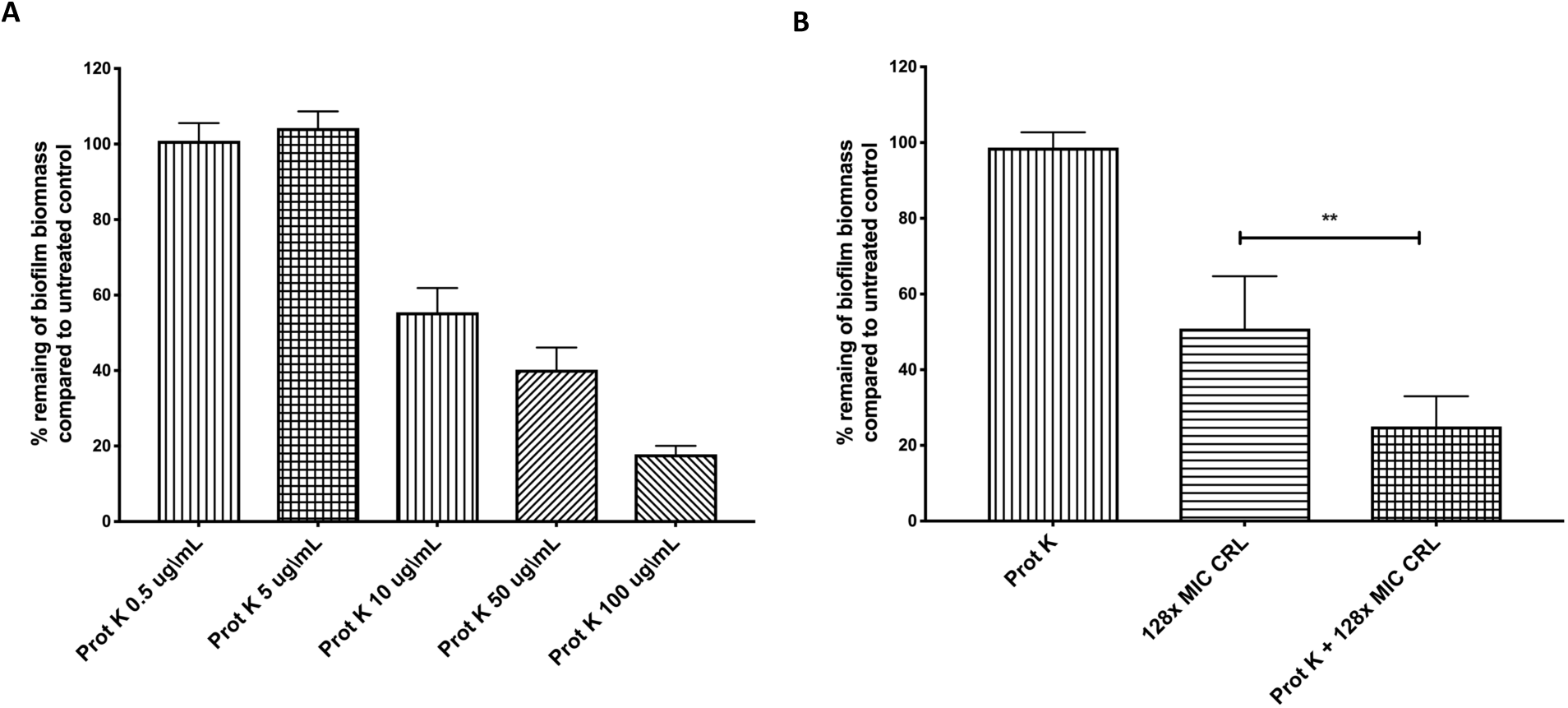
*H. pylori* G27 pre-formed biofilms are more sensitive to combination of proteinase K and clarithromycin 128-times the MIC. **(A)** 2-day old biofilms were exposed to different concentration of proteinase K along or (B) combined with and clarithromycin for 24 hours. Results present the percentage of remaining biofilm compared to the untreated control (fresh media without antibiotic and/or proteinase K). Experiments were performed three independent times with at least 8 technical replicates for each. Error bars represent standard errors of the mean. Statistical analyses were performed using ANOVA (**, *P* <0.01)

### Distinct expression profiles between biofilm and planktonic grown cells

The work presented above suggests that *H. pylori* cells grown in biofilm conditions are phenotypically distinct from those grown as planktonic ones. Thus, we decided to identify the major transcriptomic features of G27 biofilm-grown cells using an RNA-sequencing (RNA-seq). We recently used transcriptomics to learn more about the biofilm growth state of the mouse-infecting *H. pylori* strain SS1 (Hathroubi, Zerebinski et al. 2018), so decided to use the same approach to characterize the strong biofilm-producer strain G27. Initially we hoped to analyze biofilm and planktonic cells grown in the same well, but we could not because the presence of the thin pellicle prevented us from getting a clean separation of planktonic and biofilm cells. We thus decided to grow the populations separately. We assessed global patterns of transcriptomes between cells grown in a biofilm condition in 6 well dishes versus a planktonic liquid shaking growth condition. Cells from three biological replicates (two technical replicates) of each mode of growth were collected and subjected to RNA extraction. A total of 9-16 million reads per sample were generated by RNA-seq and mapped to *H. pylori* G27 complete reference genome (Baltrus, Amieva et al. 2009). These reads revealed a clustering of the biofilm-grown cells in a distinct population compared to those grown in planktonic conditions (Fig. 6A). The biofilm population was more variable compared to the planktonic one, suggesting heterogenicity of biofilm sub-populations. Indeed, as mentioned above, biofilm growth is characterized by cells attached on plastic surfaces, cells at the air-liquid interface, and cells composing the pellicle. Transcriptomic analysis showed that a total of 136 of 1534 genes (8.86%) were significantly differentially expressed (*p*< 0.05 and log_2_-fold change >1 or <−1) between *H. pylori* growing in a biofilm growth mode compared to the planktonic one (Fig. 6B). Those data are relatively similar to those found in a previous *H. pylori* biofilm transcriptomic study in strain SS1 (Hathroubi, Zerebinski et al. 2018) where 8.18% of genes were significantly differentially expressed between surface attached-cells and planktonic populations in the same wells. In the current study, we found that 93 genes were significantly upregulated in cells growing in biofilm condition while 43 genes were significantly downregulated (Fig 6B, Table 2 and 3). To confirm the results obtained by RNA-seq, the relative expression of six selected genes were quantified by quantitative reverse transcription-PCR (qRT-PCR). As demonstrated in Fig S2, the same trend was observed for all measurements (Fig. S2).

**Figure 6.**
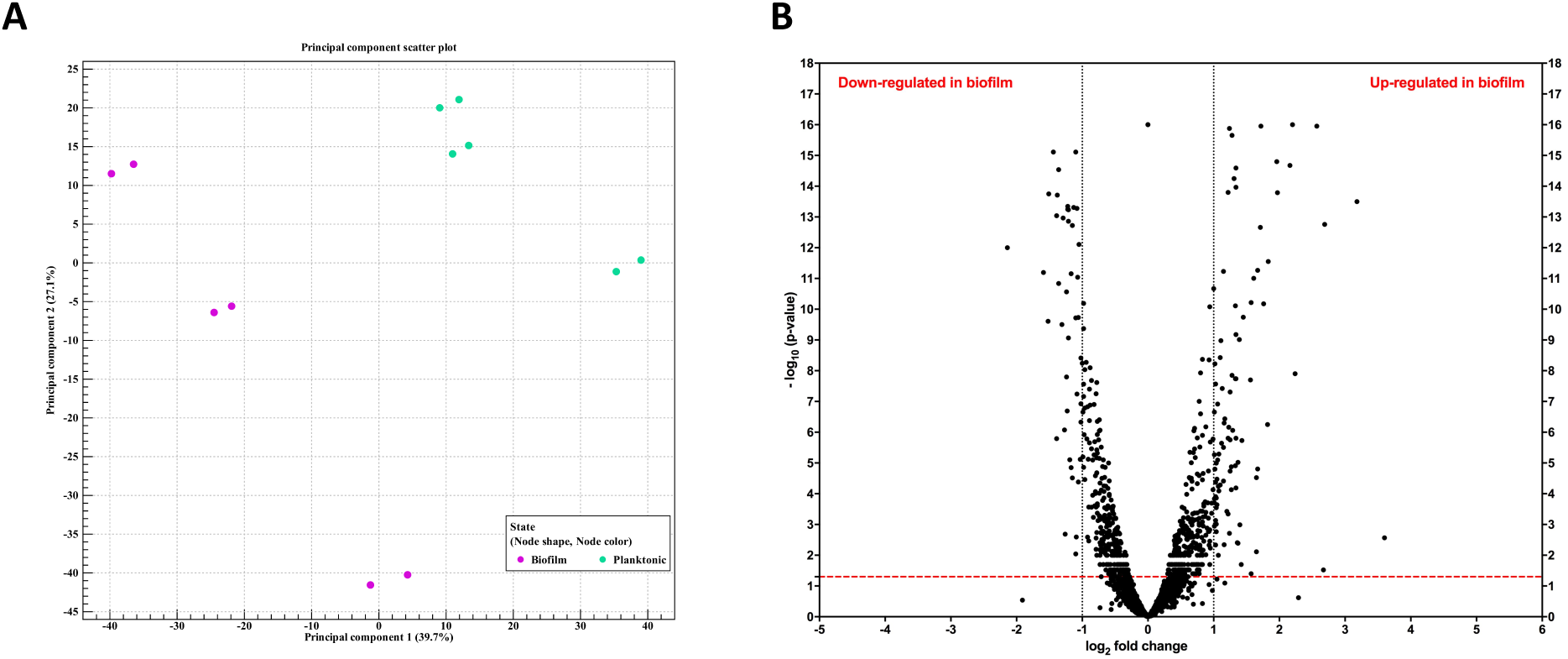
Differential gene expression in *H. pylori* biofilm-grown cells as opposed to planktonic grown cells. *H. pylori* strain G27 was grown as a biofilm or planktonic culture for 3 days, after which RNA was collected and sequenced. **(A)** Principal component analysis (PCA) of gene expression obtained by RNA-seq between biofilm growing cells (n =3) and planktonic ones (n =3) **(B)** Volcano plot of gene expression data. The y-axis is the negative log10 of P-values (a higher value indicates greater significance) and the x-axis is log2 fold change or the difference in abundance between two population (positive values represent the up-regulated genes in biofilm and negative values represent down-regulated genes). The dashed red line shows where *P* =0.01, with points above the line having *P* < 0.01 and points below the line having *P* > 0.01.

**Table 2.**
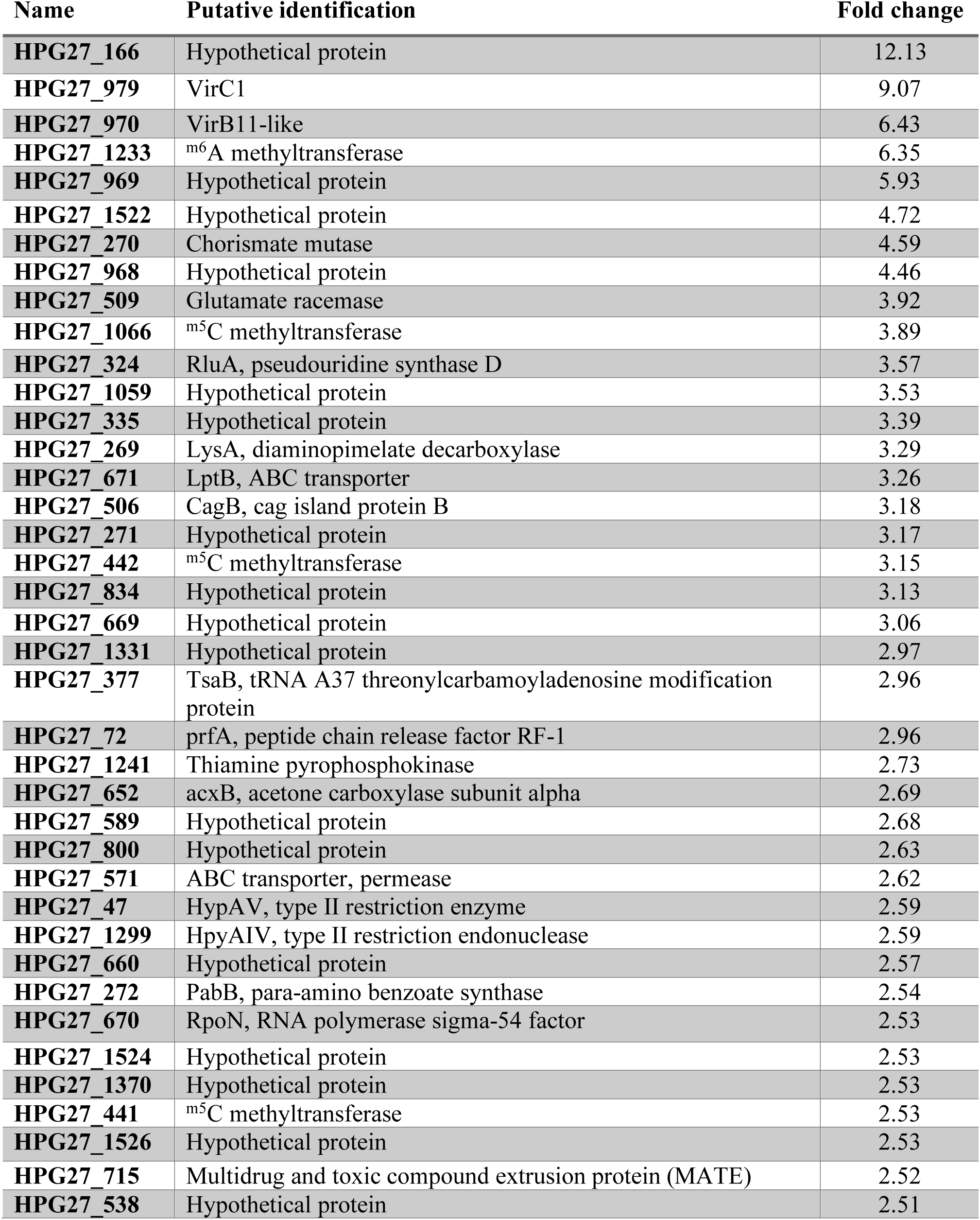
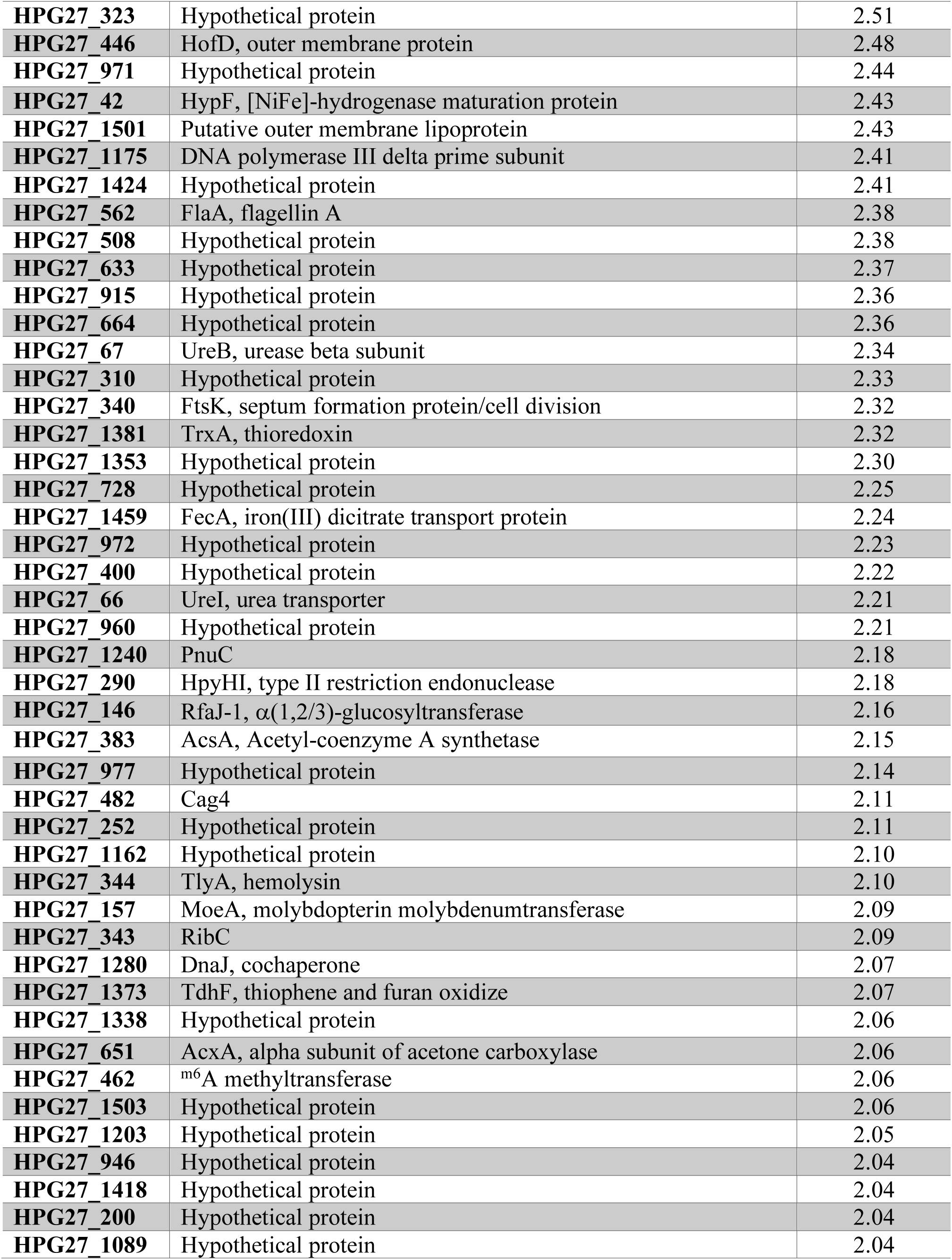
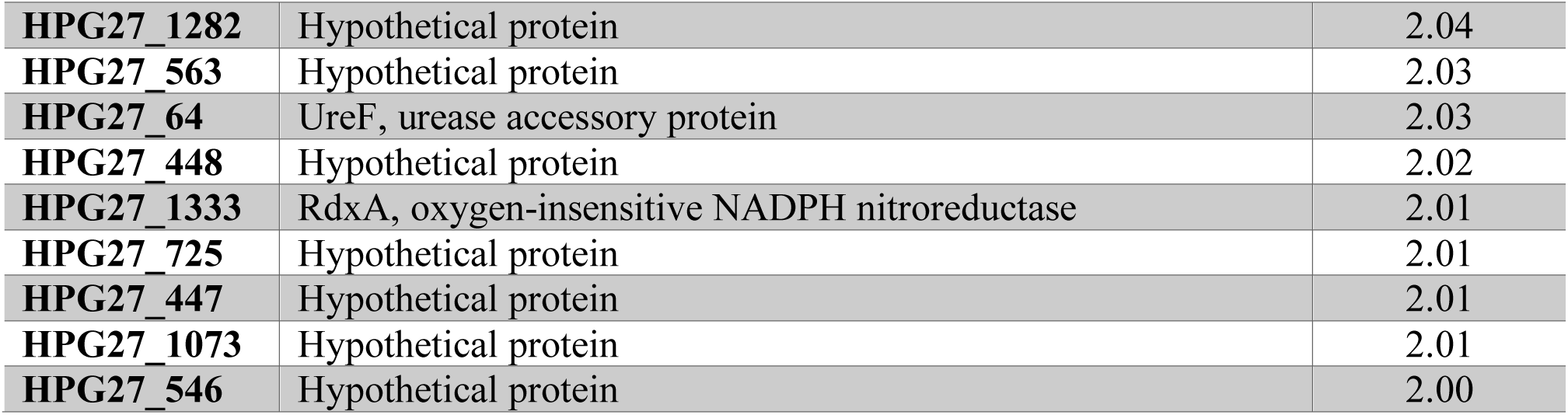
Up-regulated gene in H. pylori G27 grown in biofilm condition (cutoff ratio ≥ 1 log2 fold change and p-value <0.05) using RNA-seq analysis.

**Table 3.**
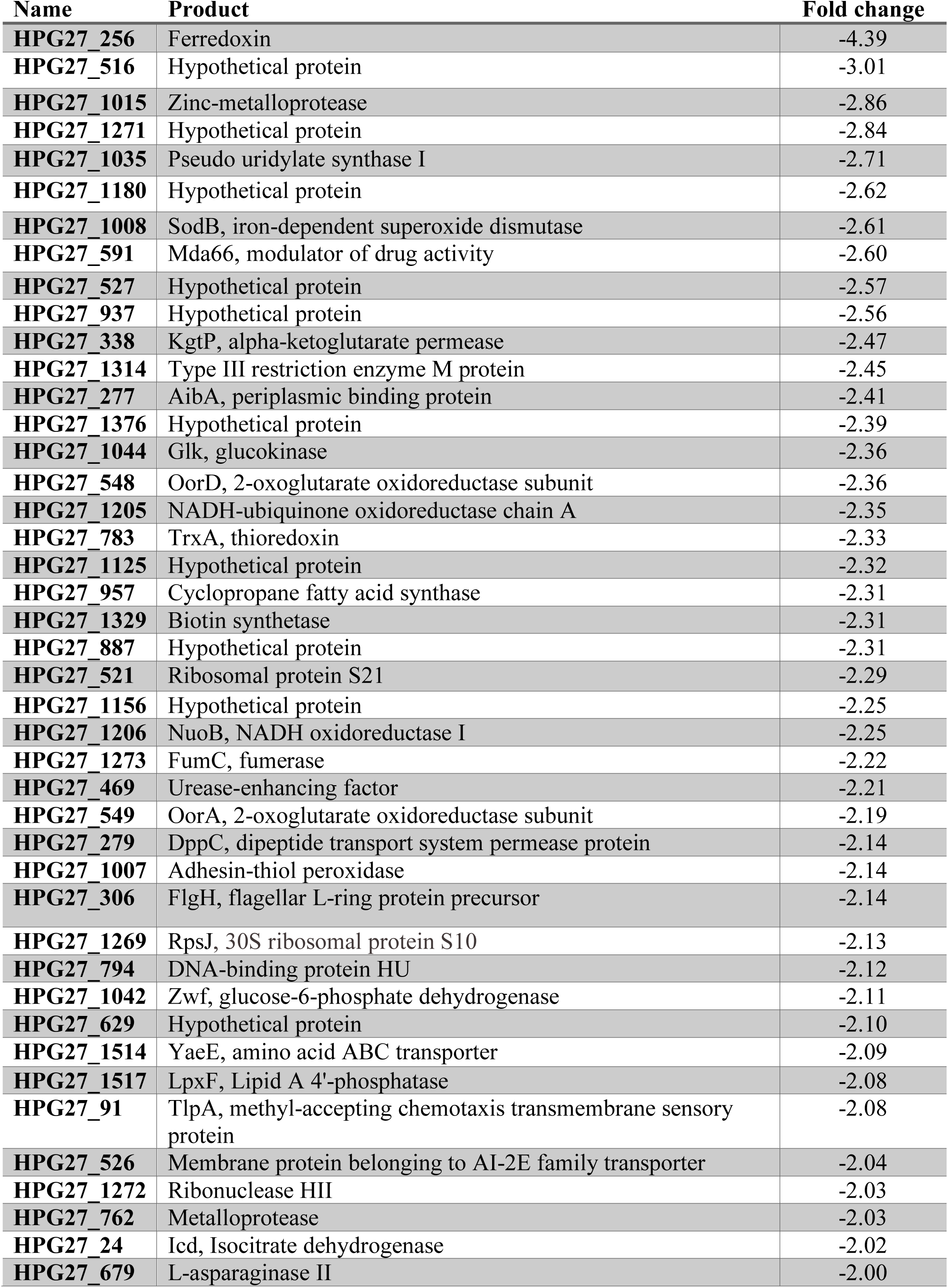
Down-regulated genes in H. pylori G27 grown in planktonic condition (cutoff ratio ≥ 1 log2 fold change and p-value <0.05) using RNA-seq analysis, grouped by functional role categories.

In overview, we found several genes belonging to T4SS systems up-regulated in cells grown as biofilms as well as genes encoding for restriction restriction-modification (R-M) and DNA repair and protection. Metabolism and related genes belonging to tricarboxylic acid (TCA) cycle were down regulated in biofilm grown cells but transcript levels for genes encoding hydrogenase, urease, and acetone carboxylase were all up (Table 2). Several genes related to the flagellar apparatus and its regulation were highly up-regulated in biofilm cells (i.e. *rpoN* and *flaA*). *rpoN* encodes for sigma 54, an important regulator for the expression of class II flagellar genes (Niehus, Gressmann et al. 2004) while *flaA* encodes the major flagellin. Flagellar filaments have previously been demonstrated to play a role in biofilm formation of both *H. pylori* strains SS1 and G27 (Hathroubi, Zerebinski et al. 2018), and thus it is not surprising to find flagellar related genes upregulated in biofilms. Additionally, *lptB*, which encodes for a lipopolysaccharide export system ATPase, was up-regulated. This gene was also previously up-regulated in biofilm cells of *H. pylori* strain SS1 (Hathroubi, Zerebinski et al. 2018).

Among the down-regulated genes in cells grown in biofilm conditions, we found two genes related to quorum sensing, HPG27_277 and HPG27_526, which encode for the AI-2 periplasmic binding protein A (AibA) and a putative AI-2 exporter, respectively. AI-2 is a quorum-sensing molecule produced by the metabolic enzyme LuxS and a chemorepellent molecule senses by *H. pylori* (Rader, Wreden et al. 2011, Anderson, Huang et al. 2015). Previous studies have shown that a *luxS*-deficient of aibA-deficient *H. pylori* mutant exhibited increased levels of adherence and biofilm (Cole, Harwood et al. 2004, Anderson, Huang et al. 2015) and reciprocally the *luxS* overproducer strain (overproduction of AI-2) induced a dispersion of biofilm cells (Anderson, Huang et al. 2015). In this study, the gene encoding for LuxS was significantly down-regulated (−1.26 fold-change, p<0.05) but below our cutoff of −2-fold change. Our data thus support the idea that AI-2 is important for cell dispersion in biofilms and AI-2 production, export and sensing are all downregulated during biofilm growth

Another gene downregulated in biofilm cells, *lpxF*, encodes for an enzyme responsible for dephosphorylation of the lipid A 4’-phosphate group in *H. pylori*. The LpxF phosphatase has been previously associated with biofilm formation in *H. pylori* and inactivation of its gene resulted in increased biofilm formation and bacterial fitness (Gaddy, Radin et al. 2015). Together, these results strongly support that our transcriptomic approach has identified key biofilm genes; several additional ones are discussed in the discussion section.

### Identification and characterization of biofilm mutants

As an alternative method to identify genes involved in *H. pylori* biofilm formation, we screened an *H. pylori Tn*-7 based transposon-based mutant library (Salama, Shepherd et al. 2004) for biofilm-defective mutants. The pool of mutants forms biofilms on polystyrene plates, and we thus embarked on a selection for mutants unable to form biofilms. After incubation, the supernatant, which contains non-attached planktonic cells and potential-biofilm defective mutants, was transferred to a new sterile plate every 3 hours for the first 12 hours and every 12 hours for the next 72 hours. The goal of this approach was to enrich our supernatant with potential biofilm-defective mutants. At the end of the 72-hour incubation period, supernatant cells were plated and 97 colonies were randomly picked and assessed for biofilm formation using crystal violet staining. Eight of these displayed significant (*p*<0.05 or *p*<0.01) biofilm formation defects. The eight biofilm-defective mutants were re-tested and confirmed to form significantly less biofilm than the parental strain (*p*<0.05 or *p*<0.01) over of six days of growth (**Fig. 7**).

**Figure 7.**
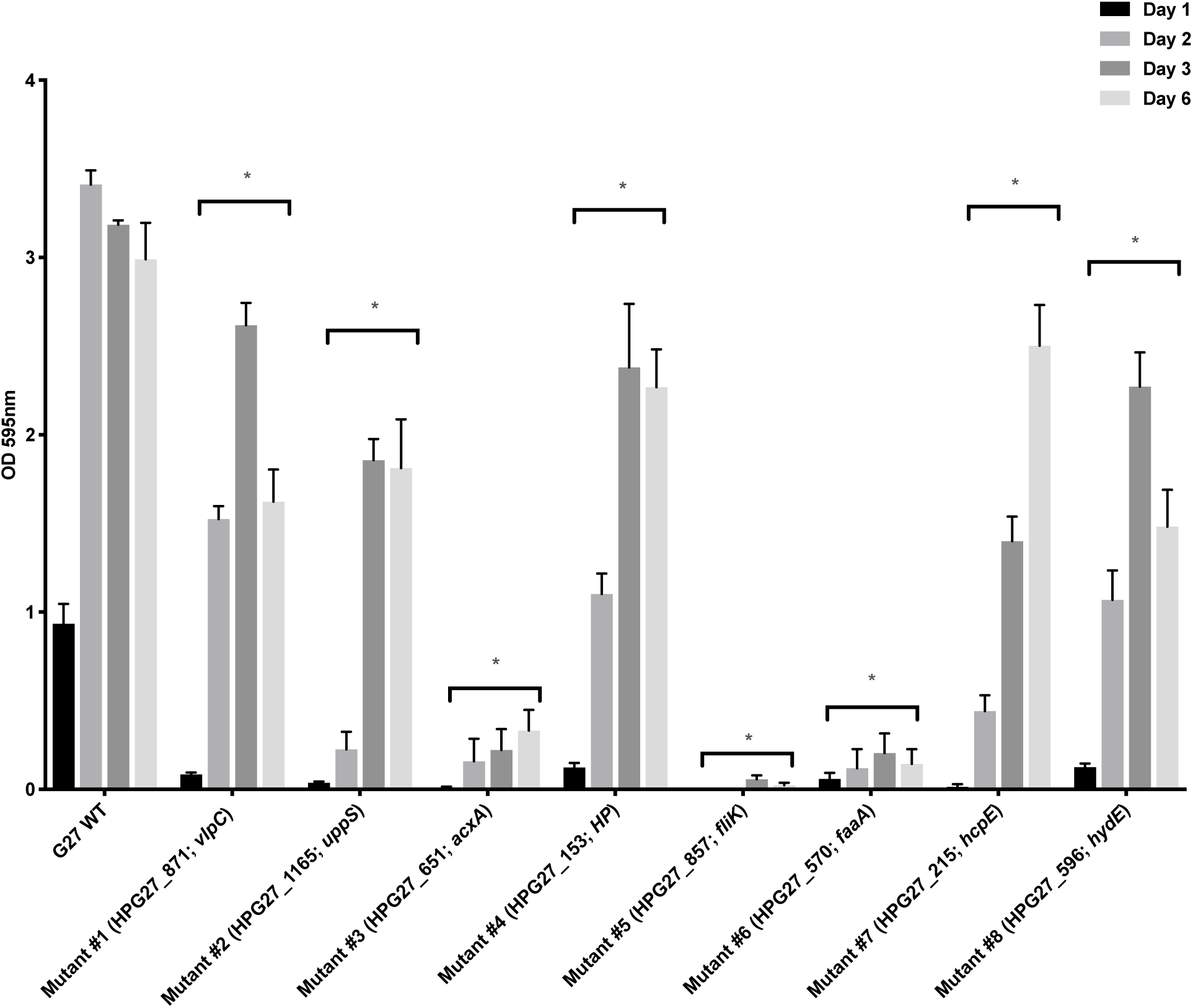
Biofilm formation by *H. pylori* G27 WT and biofilm-defective mutants. Biofilm formation was assessed using the microtiter plate biofilm assay over three days. Results represent the crystal violet absorbance at 595nm which reflect the biofilm biomass. Experiments were performed three independent times with at least 3 technical replicates for each. Error bars represent standard errors for each average value. HP; Hypothetical protein, WT; Wild-type. Statistical analyses were performed using ANOVA (*, *P*<0.05)

To further confirm the biofilm defects, these mutants were analyzed by CSLM (**Fig. S4**). As shown in Fig. S4, biofilm defective-mutants were all affected in forming biofilm compared to wild-type parental strain G27. The mutants with the strongest defect in the CV assay, mutants #3, #5 and #6, displayed only some dispatched microcolonies and were severally impaired in their ability to form a developed biofilm (**Fig. S4**). To ensure that biofilm defect was not associated to a defect in growth, we measured the growth rate and compared it to the wild-type strain (data not shown). From the eight mutants, only mutant #2 demonstrated a slight decrease in its growth rate.

Using nested PCR and a combination of transposon-specific primers, we created PCR products for each transposition events. Sequencing of these PCR products identified eight non-redundant disrupted genes (**Fig. S5**), listed in Table 4. Two of the mutants had insertions in genes encoding for vacuolating toxin A (VacA)-like proteins, *vlpC* (mutant #1) and *faaA* (mutant #6). Those genes are among the largest genes in the *H. pylori* genome and encode membrane proteins (Radin, Gaddy et al. 2013). Interestingly, FaaA localizes to the flagellar sheath, and is required for proper flagella function and for mouse colonization (Radin, Gaddy et al. 2013). We identified another insertion in gene related to flagella, HPG27_857 (mutant #5) which encodes for a flagella hook-length control protein (FliK). From 8 identified mutants two belonged to flagellar-associated proteins, which again supports the tight connection between *H. pylori* biofilm and flagella.

**Table 4.** H. pylori G27 reduced-biofilm mutants identified and sequenced in the present study.

| **Mutant ID** | **NCBI Locus tag** | ***Tn* location (bp)** | **Description of gene** | **Functional class** |
| --- | --- | --- | --- | --- |
| **#1** | **HPG27_871** | 952330 | VlpC, VacA-like protein, | Virulence |
| **#2** | **HPG27_1165** | 1284748 | UppS, putative undecaprenyl pyrophosphate synthase<br>Isoprenyl transferase | Peptidoglycan synthesis |
| **#3** | **HPG27_651<sup>♥</sup>#** | 712390 | AcxA, alpha subunit of acetone carboxylase | Virulence/colonization/motility |
| **#4** | **HPG27_153** | 171244 | Hypothetical protein | Unknown function |
| **#5** | **HPG27_857** | 922300 | FliK, flagellar hook-length control protein | Motility |
| **#6** | **HPG27_570** | 622795 | FaaA, VacA-like protein | Motility/Virulence/colonization |
| **#7** | **HPG27_215** | 241131 | HcpE, cystein riche protein | Virulence |
| **#8** | **HPG27_596<sup>#</sup>** | 650623 | HydE | Hydrogenase synthesis |

<sup>♥</sup>HPG27\_651 gene was also found up-regulated in biofilm cells
<sup>#</sup>Gene knockout constructs, *AcxAB::Cm* and *hydE::Cm* were generated to confirm the observed phenotype

Interestingly, one insertion was in the gene *acxA* (HPG27_651; mutant #3), which encodes the alpha subunit of acetone carboxylase. The same gene was found up-regulated in biofilm cells (Table 2), reinforcing the idea that this enzyme may play role in *H. pylori* biofilm formation. Sequencing also revealed an insertion in *hydE*, the last of gene of hydrogenase operon *hydABCDE*, involved in localizing the hydrogenase complex to the membrane. Mutation in *hydE* gene was previously demonstrated to abolish hydrogenase activity in *H. pylori* (Benoit, Mehta et al. 2004).

### Genes essential for biofilm formation

Based our both approaches we made a short list of candidate genes and generated mutants in each gene as listed in Table 1. Some of these were to remake the transposon insertions with targeted deletion/insertions, and others were to test the importance of upregulated genes from the transcriptomic data. The loss of these genes did not affect growth except for *ΔacsA* which demonstrated a slight delay in growth which may impact on its ability to make biofilm. As shown in Figure 8, biofilm formation ability was evaluated. All mutants demonstrated substantially reduced biofilm formation (Fig. 8). During our *H. pylori Tn*-7 mutant library screen, we found that insertions in *hydE* and *acxA* decreased biofilm formation (Fig, 7). Deletion of *hydE* and *acxAB* genes showed a decrease biofilm formation (> 11-fold and > 2.5-fold, respectively), supporting the idea that these genes play an important role in *H. pylori* biofilm formation (Fig. 8).

**Figure 8.**
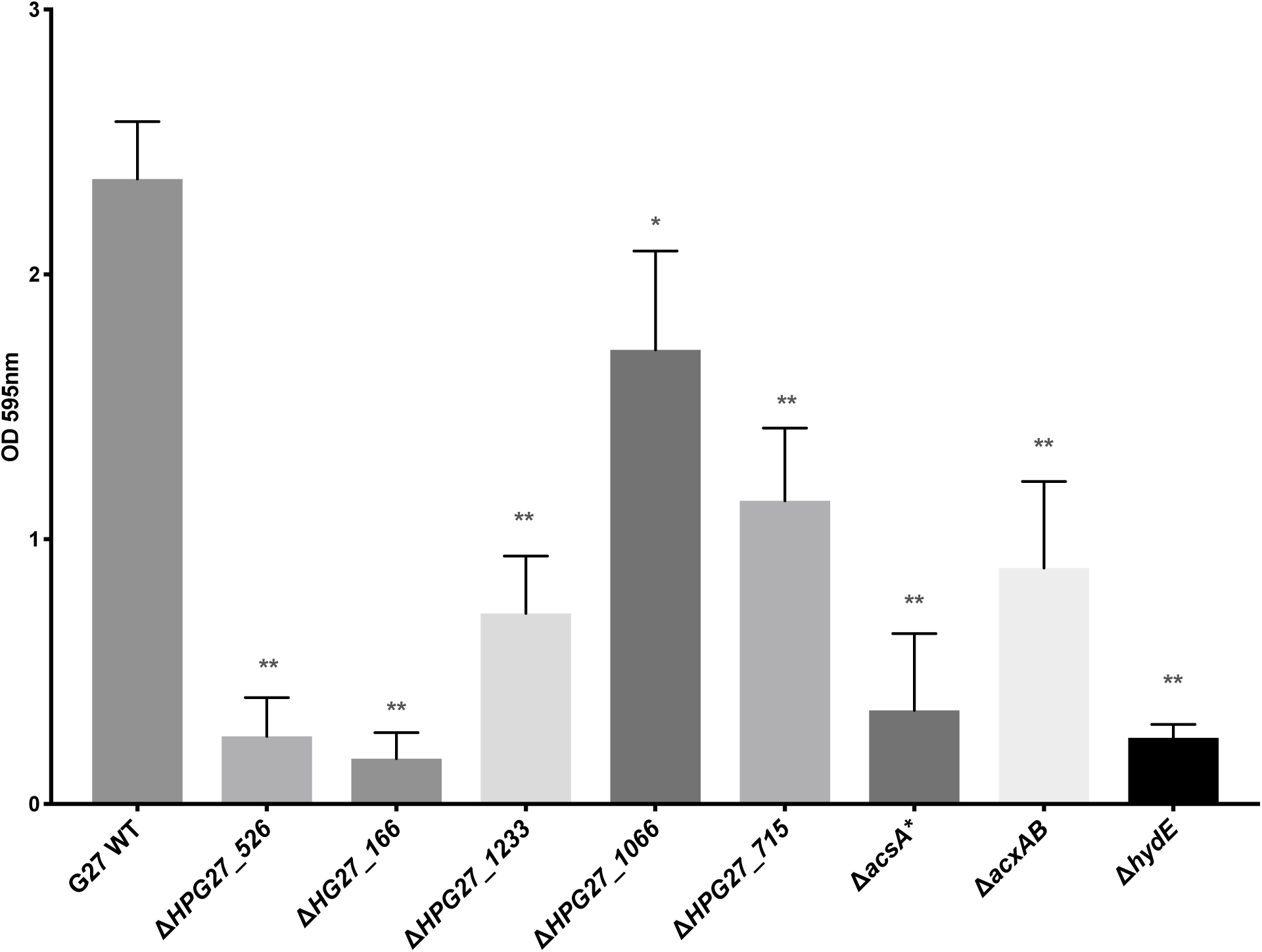
Biofilm formation by *H. pylori* G27 WT and its isogenic mutants. Biofilm formation was assessed using the microtiter plate biofilm assay after three days of growth. Results represent the crystal violet absorbance at 595nm which reflect the biofilm biomass. Experiments were performed two independent times with at least 6 technical replicates for each. Error bars represent standard errors for each average value. Statistical analyses were performed using ANOVA (*, *P* <0.01; **, *P* <0.0001)

In our transcriptomic analysis, HPG27_166, HPG27_1233, HPG27_1066 and HPG27_715 were all significantly upregulated during biofilm growth. Their inactivation significantly decreased biofilm formation in *H pylori* G27 (*P* <0.01 or *P* <0.0001) (Fig. 8). Deletion of gene HPG27_166, which demonstrated the highest expression level during biofilm growth, resulted in a complete loss of biofilm formation. HPG27_1066 as well as HPG27_1233 encode for methyltransferases they well known to contribute to changes of the epigenome (Jeltsch 2002). HPG27_1233, called JHP1050 in *H. pylori* J99, was recently demonstrated to affect the transcription of multiple genes, adherence to host cell, natural competence and bacterial cells shape (Estibariz, Overmann et al. 2019). Here we demonstrated that HPG27_1233 also affects biofilm growth and its inactivation results in a significant decrease in biofilm formation of *H. pylori* G27. Inactivation of HPG27_526 gene, encoding the putative AI-2 transporter, surprisingly resulted in severe defect in biofilm formation. This gene was downregulated in the biofilm transcriptomics, so we predicted that loss would result in elevated biofilm. However, this gene product has yet to be characterized in *H. pylori*, so it may play a different role, or AI-2 may play more nuanced role in biofilm formation.

## DISCUSSION

We report here a characterization of biofilm formation in *H. pylori* G27, a strain that forms robust biofilms under many conditions. *H. pylori* G27 biofilm cells exhibited enhanced tolerance to antibiotics compared to planktonic-grown cells. Treating with proteinase K, which targets extracellular proteins including the protein-based biofilm matrix, significantly increased clarithromycin efficacy. Using two different approaches, transcriptomic and mutant identification, we revealed many genetic determinants for biofilm growth including a set of genes that suggests flagella production, slow growth, and altered metabolism are keys during this mode of growth.

Biofilm formation appears to be a common trait among *H. pylori* strains but has been largely unexplored. Contributing to the lack of studies is the fastidious nature of *H. pylori* as well as a clear lack of consensus among the techniques and model strains to be used to study biofilms in *H. pylori*. In this study, we worked with *H. pylori* G27, which is a strong biofilm former when grown under standard *H. pylori* growth conditions, and has become a commonly studied H. pylori strain (Servetas, Carpenter et al. 2016, Hathroubi, Zerebinski et al. 2018, Windham, Servetas et al. 2018). *H. pylori* G27 appears to be a good model for biofilm studies because it exhibits a reproducible 3-dimensional and dense structure containing communities of cells tightly packed and connected to each other and to the surface, with pili-like structures and abundant flagellar filaments.

Given the visible detection of flagella in the biofilm (Hathroubi, Zerebinski et al. 2018) it is not surprising to find genes encoding for flagella expression in the transcriptomic of biofilms (i.e. *rpoN* and *flaA*) as well as during our screen for biofilm-defective mutants (*faaA* and *fliK*). These findings reinforced the idea that was recently developed that flagella can play an important structural role in biofilms in *H. pylori* and in *E. coli* (Serra, Richter et al. 2013, Hathroubi, Zerebinski et al. 2018, Rizzato, Torres et al. 2019).

Biofilm cells were phenotypically distinguishable from those grown as planktonic ones and exhibit marked tolerance to antibiotic killing, consistent with previous results using distinct *H. pylori* strains (Yonezawa, Osaki et al. 2013, Yonezawa, Osaki et al. 2019). We expanded this initial work to show that this phenotype may be related to the biofilm matrix-associated proteins. These proteins were previously described to contribute to the structure and stability of *H. pylori* biofilm (Hathroubi, Zerebinski et al. 2018, Windham, Servetas et al. 2018). When proteinase K is used in combination with clarithromycin, a synergistic effect on antibiotic efficacy was observed. This outcome suggests that proteins may act as a protective shield and support the idea that weakening the matrix-based proteins enhances the efficiency of clarithromycin treatment. However, combined treatment was not enough to eradicate the pre-formed biofilm suggesting other mechanisms to play a role such as low metabolism (Hathroubi, Zerebinski et al. 2018).

In this work, we demonstrated that biofilm cells displayed a distinct transcriptome. Several genes related to the cell envelope were up-regulated in the biofilm, including many uncharacterized membrane and outer membrane proteins (i.e. HPG27_446, HPG27_571, HPG27_1370, HPG27_1501 and HPG27_715) (Table 2). Several of these already stand out as interesting proteins. HPG27_715 encodes for a multi drug and toxic compound extrusion (MATE)-type multidrug efflux pump (Chen, Ye et al. 2018). Inactivation of gene HPG27_715 significantly decreased biofilm formation. However, its role in antimicrobial tolerance during biofilm formation in *H. pylori* has yet to be determined.

Among the other genes up-regulated during biofilm growth we found the *lptB* gene, which encodes for a lipopolysaccharide export system ATPase. This gene has been previously reported to be up-regulated in the biofilm of *H. pylori* strain SS1 (Hathroubi, Zerebinski et al. 2018). Another gene related to LPS, *rfaJ-1* was also found to be up-regulated in biofilm cells. This gene encodes for an *α*-1,6-glucosyltranferase that plays an integral role in the biosynthesis of the core LPS (Logan, Altman et al. 2005) and the loss of this gene resulted in a truncated LPS and a defect in mouse colonization (Altman, Chandan et al. 2008). These results suggest that LPS is key to *H. pylori* biofilm formation and may play a role in promoting cell-cell interactions.

Additional upregulated genes map to two of the four *H. pylori* type IV secretion systems (T4SS) (Chang, Shaffer et al. 2018). These four are the cytotoxin-associated gene pathogenicity island (*cag*PAI)-encoded T4SS (*cag* T4SS) which translocates CagA into the host gastric epithelial cells; the *comB* T4SS that mediates DNA uptake from the extracellular environment; and two poorly-characterized T4SS, *tfs3* and *tfs4*, which both have been suggested to play a role in horizontal DNA transfer (Yuan, Wang et al. 2018). In this study, we found several genes belonging to the *cag* and *tsf4* T4SS systems up-regulated in cells grown as biofilms. Specifically, of the *cag* T4SS, upregulated genes were *cagB*, encoding one of the Cag ATPases, and *cag4*/*cagγ*, encoding a cell wall hydrolase of the *cag* T4SS. Of the *tfs4* system, many genes including i.e *virC1*, *virD4, virb11-like*, HPG27_977, HPG27_972, HPG27_971, HPG27_969 and HPG27_968 (see Fig S3). The pili-like structures observed using SEM may belong to this *tfs4* system however, further investigations are needed to confirm their exact origin. The role of *tfs4* during biofilm formation is not yet understood but it could increase DNA transfer between biofilm cells. This idea is supported by recent data from *Bacillus subtilis*, where biofilm formation drove high rates of conjugative ICE transfer compared to planktonic cells (Lecuyer, Bourassa et al. 2018). Such a situation might also increase the requirement for restriction-modification (R-M) systems, of which several were upregulated (Table 2) or required for biofilm formation (Fig. 8).

Biofilm cells also displayed pronounced metabolic changes including down-regulation of TCA cycle activity, along with increased activity of both acetone carboxylase and hydrogenase. Downregulated tricarboxylic acid (TCA) cycle enzymes and electron transport chain components include glucokinase (*glk*), citrate synthase (*gltA*), isocitrate dehydrogenase (*icd*,), fumarase (*fumC*), NADH-ubiquinone oxidoreductase (*nqo10*) and NADH oxidoreductase I (*nuoB*). These data suggest that cells grown in biofilm may display decreased flux through the TCA cycle as either a way to overall lower metabolism or alter it. Similar outcomes were observed previously during biofilm formation in strain SS1 (Hathroubi, Zerebinski et al. 2018). This lowered growth state can also contribute to antibiotic tolerance as described for other organisms (Stewart 2015, Ciofu and Tolker-Nielsen 2019).

Not all metabolic genes were low, indeed, HPG27_651 and HPG27_652 genes, located in the acetone carboxylase operon HG27_651-653 (*acxABC*) were significantly upregulated in cells gown in biofilm condition (Table 3). This operon encodes enzymes implicated in the ATP-dependent carboxylation of acetone to acetoacetate (Brahmachary, Wang et al. 2008). Acetoacetate can be then converted acetoacetyl-CoA, which can be metabolized further to generate two molecules of acetyl-CoA (Brahmachary, Wang et al. 2008). Our mutant screen also found one biofilm-defective mutant with an insertion in gene HPG27_651 (*acxA*), which lends further support to the importance of this process in *H. pylori* biofilm growth. To confirm the role of acetone carboxylase in biofilm formation of *H. pylori*, we inactivated *acxAB* and evaluated its biofilm formation. As expected, *ΔacxAB* mutant showed a loss of ability to form a biofilm which suggest a role of acetone carboxylase during this mode of growth.

Another route to produce acetyl-CoA was also upregulated in biofilm growth, *acsA*. This gene encodes the AcsA acetyl-coenzyme A synthetase thatgenerates acetyl CoA through the metabolism of acetate. The acetyl-CoA pathway begins with the reduction of a carbon dioxide to carbon monoxide in *H. pylori* via the AcsA (Kuhns, Benoit et al. 2016). The amount of this synthetase (expression and activity) is augmented when cells are using molecular hydrogen, as an electron donor (Kuhns, Benoit et al. 2016). One idea is that during biofilm formation, acetone may be used as an alternative carbon source, although it is not yet clear where the acetone comes from. It seems reasonable to predict that the biofilm cells would turn to an unusual carbon source in order to retain respiratory metabolism under an otherwise carbon-poor nutrient condition. Inactivation of this gene in *H. pylori* significantly decreased biofilm formation, however, this mutant showed also a slight defect in growth which may be associated with the effect in biofilm formation.

Interestingly, we also found that one gene encoding for part of the hydrogen-oxidizing hydrogenase, HypF, was upregulated. HypF is [NiFe]-hydrogenase maturation protein F and has been implicated in the maturation of nickel-iron containing enzymes including hydrogenase in *H. pylori* (Olson, Mehta et al. 2001, Benoit, Mehta et al. 2004). Hydrogen-oxidizing hydrogenases catalyze the reversible oxidation of molecular hydrogen to provide cells with a high-energy electron donor for respiration-based energy metabolism (Olson and Maier 2002) (Kuhns, Benoit et al. 2016). Hydrogenase is thought to be important when carbon sources are in short supply. In support of the importance of hydrogen metabolism in biofilm cells, we found a biofilm-defective mutant with an insertion in *hydE*, the last gene of the hydrogenase operon *hydABCDE* (Benoit, Mehta et al. 2004). In order to validate the importance of *hydE* gene, we generated a *ΔhydE* mutant and demonstrated that the loss of *hydE* lead to an impairment of biofilm formation. Finding these genes required for hydrogenase activity and biofilm in *H. pylori* using two different approaches suggest that oxidation of hydrogen by hydrogenase might be important for biofilm growth in *H. pylori*.

In conclusion, our findings showed that *H. pylori* is capable of adjusting its phenotype when grown as biofilm, having altered gene expression profiles, changing its metabolism and elevating gene of membrane proteins including those encoding *tfs4* and flagella. In particular, it seems that *H. pylori* shifts its metabolism to rely on molecular hydrogen and the production of acetyl Co-A. This growth state creates *H. pylori* that are highly antibiotic tolerant. It will be exciting to learn how antibiotic tolerance is created and how this of growth can be targeted to promote more effective *H. pylori* treatments.

## Acknowledgements

This work was supported by National institute of Allergy and Infectious Diseases (NIAID) grant RO1AI116946 (to K.M.O) and funds from the Santa Cruz Cancer Benefits Group (to KMO). We would like to acknowledge Robert Maier (University of Georgia) and Fitnat Yildiz (University of California, Santa Cruz) for their helpful suggestions and comments on the study. We thank Nina Salama for providing the transposon library. We thank Ben Abrams (University of California, Santa Cruz) for confocal microscope training and assistance. We also thank Tom Yuvzinsky University of California, Santa Cruz) for electron microscopy and the W. M. Keck Center for Nanoscale Optofluidics for use of the FEI Quanta 3D Dual beam microscope. Finally, we thank members of the Ottemann Lab, particularly Shuai Hu and Yasmine Elshenawi, for helpful discussions and comments on the manuscript.

---

**Figure S1.**
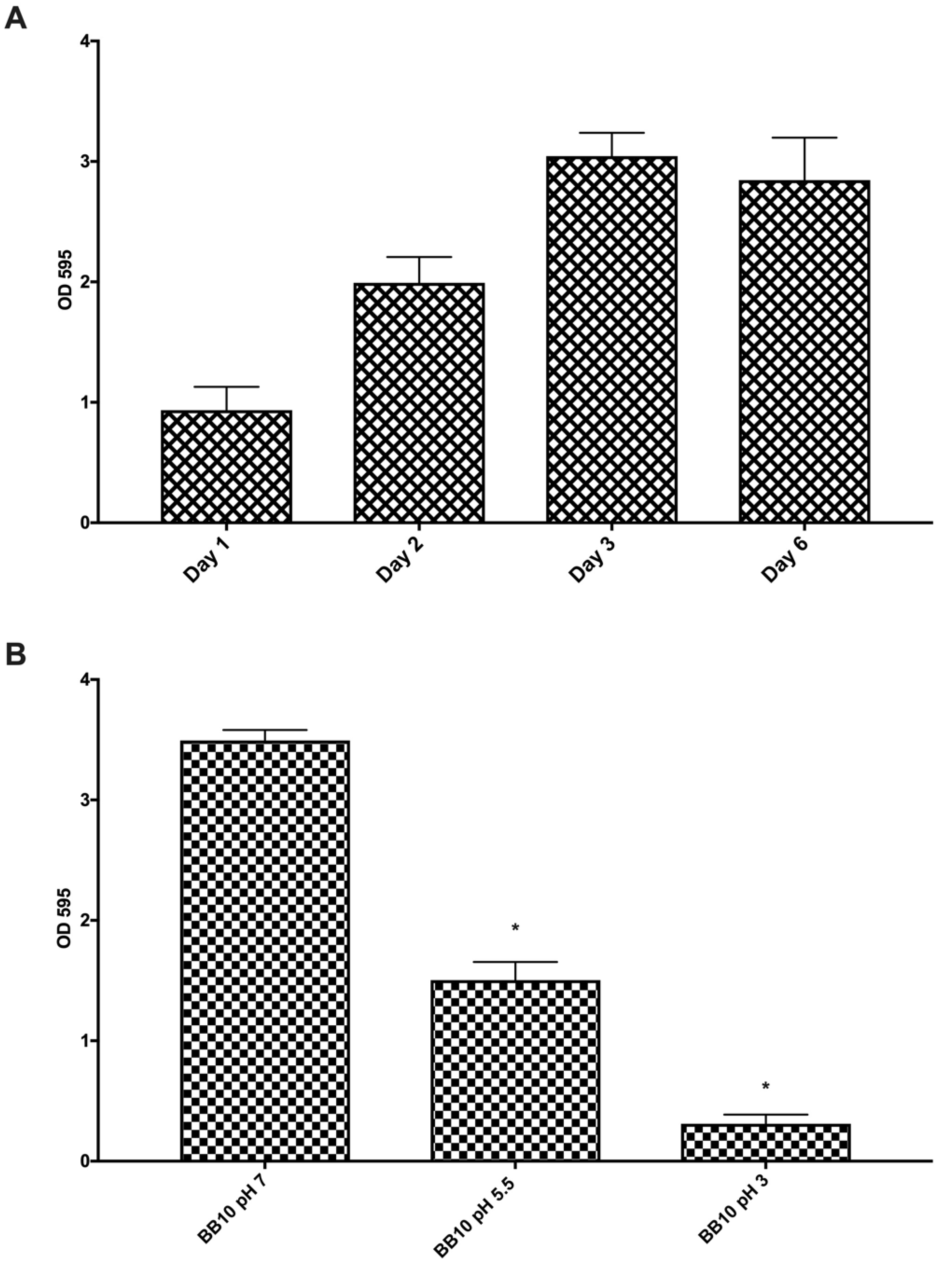
*H. pylori* G27 biofilms is time and pH-dependent. (A) Biofilm formation of *H. pylori* strains were assessed using the microtiter plate biofilm assay at different time points. (B) Effect of pH variation on biofilm development, using BB10 pHed to different values and its normal, pH 7. Results represent the crystal violet absorbance at 595nm which reflect the biofilm biomass. Experiments were performed three independent times with at least 3 technical replicates for each. Error bars represent standard errors of the mean for each average value. Statistical analyses were performed using ANOVA (*, *P* <0.05)

**Figure S2.**
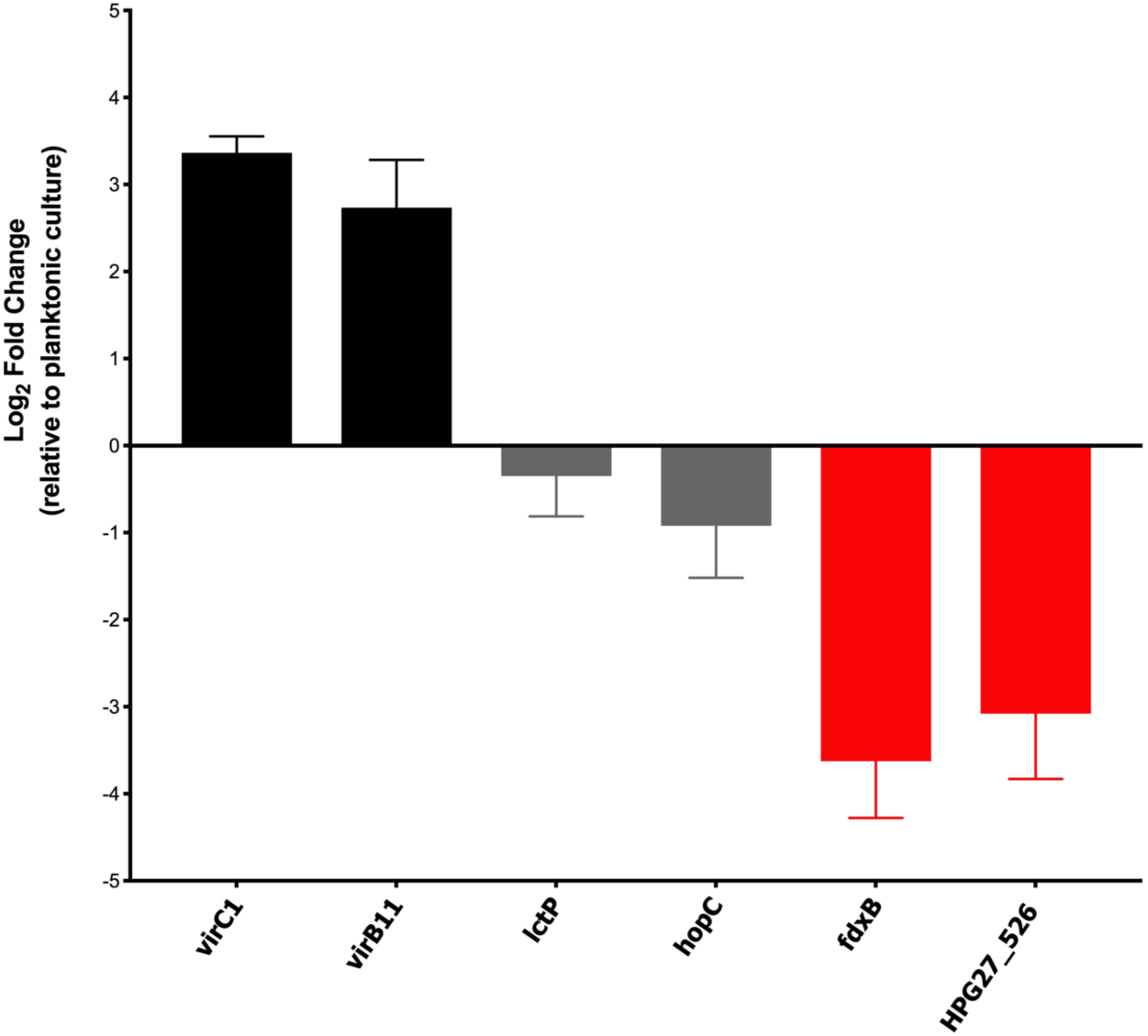
qPCR validation of the transcription of selected differentially expressed genes. The data indicate the fold change in expression of genes in H. pylori biofilm cells compared to planktonic cells. Fold changes in gene expressions were calculated after normalization of each gene with the constitutively expressed gene control *gapB*. Bars represent the mean and error bars the standard error of the mean. Black and gray bars represent qPCR and RNA-seq results, respectively. Statistical analyses were performed using threshold cycle (2^−ΔΔCT^) values, and all results were statistically significant (*P <*0.01) except for *lctP* and *hopC* gene transcripts.

**Figure S3.**
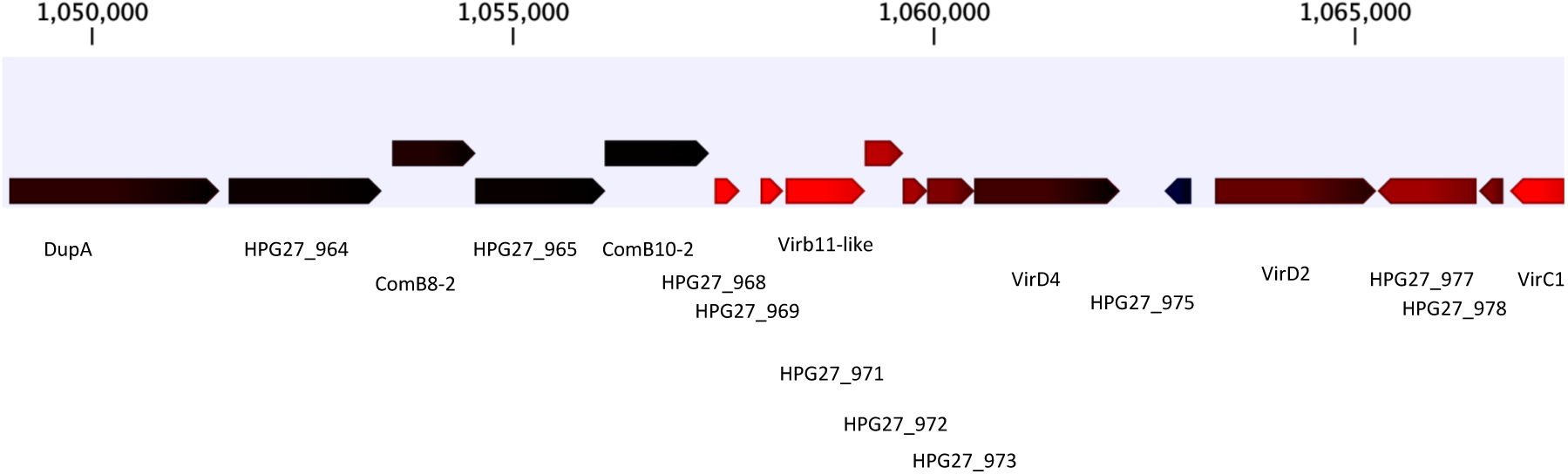
Genes expression of Type IV secretion system 4 (tfs4) genes in *H. pylori* G27 cell grown in biofilm condition. Red represent genes that are up-regulated in cells grown in biofilm condition and higher intensity of red denotes greater fold-change.

**Figure S4.**
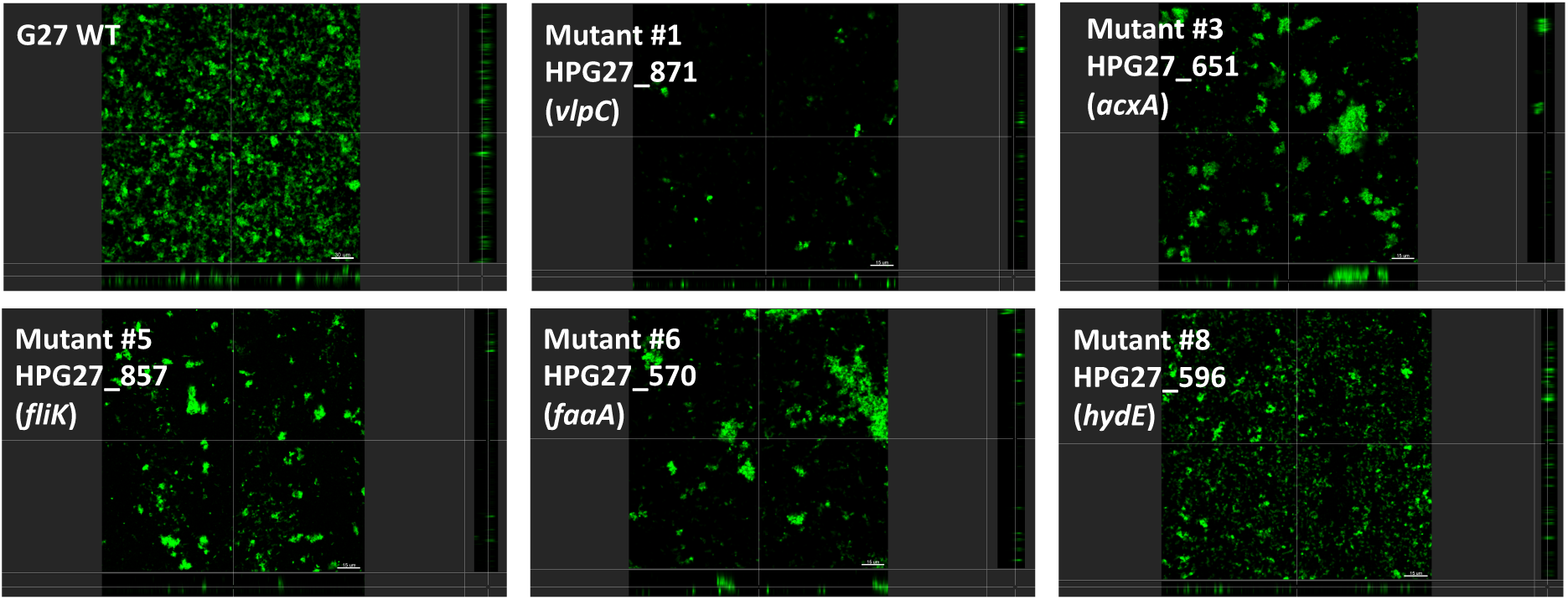
Confocal laser scanning microscopy of biofilm formation by G27 defective-mutants mutants. CLSM micrographs comparing biofilm formation of the wild type G27 strain and biofilm-defective mutants. Bacteria were grown for 3 days and then biofilm were stained with FM 1-43 which become fluorescent once inserted in the cell membrane.

**Figure S5.**
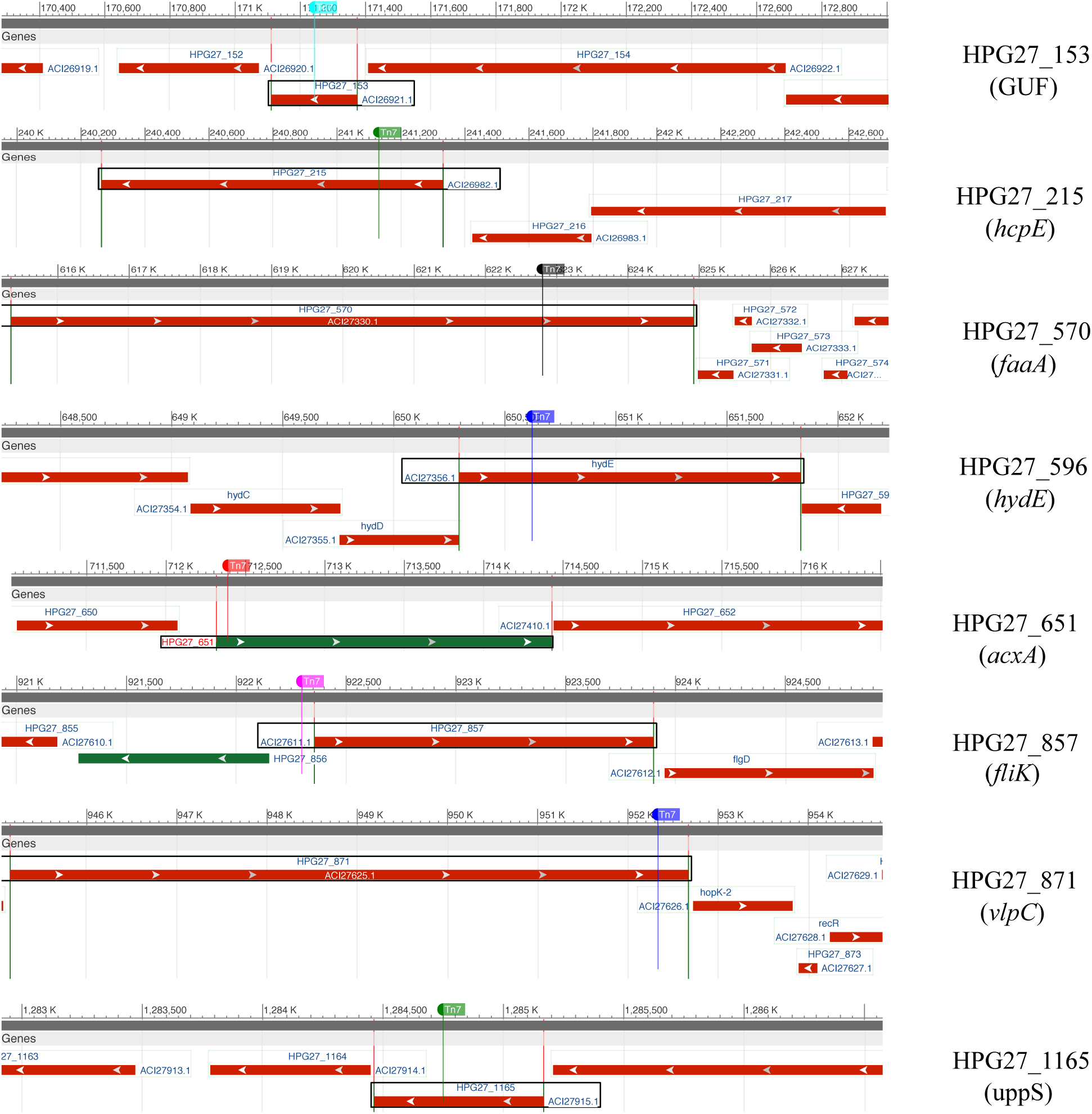
Graphical map showing the position and surrounding regions of the insertion site of Tn7 transposon in biofilm-defective mutants. Boxes represent the flanking open reading frame and colored line represent the genomic position of the transposon insertion.

**Supplemental_Table 1.**
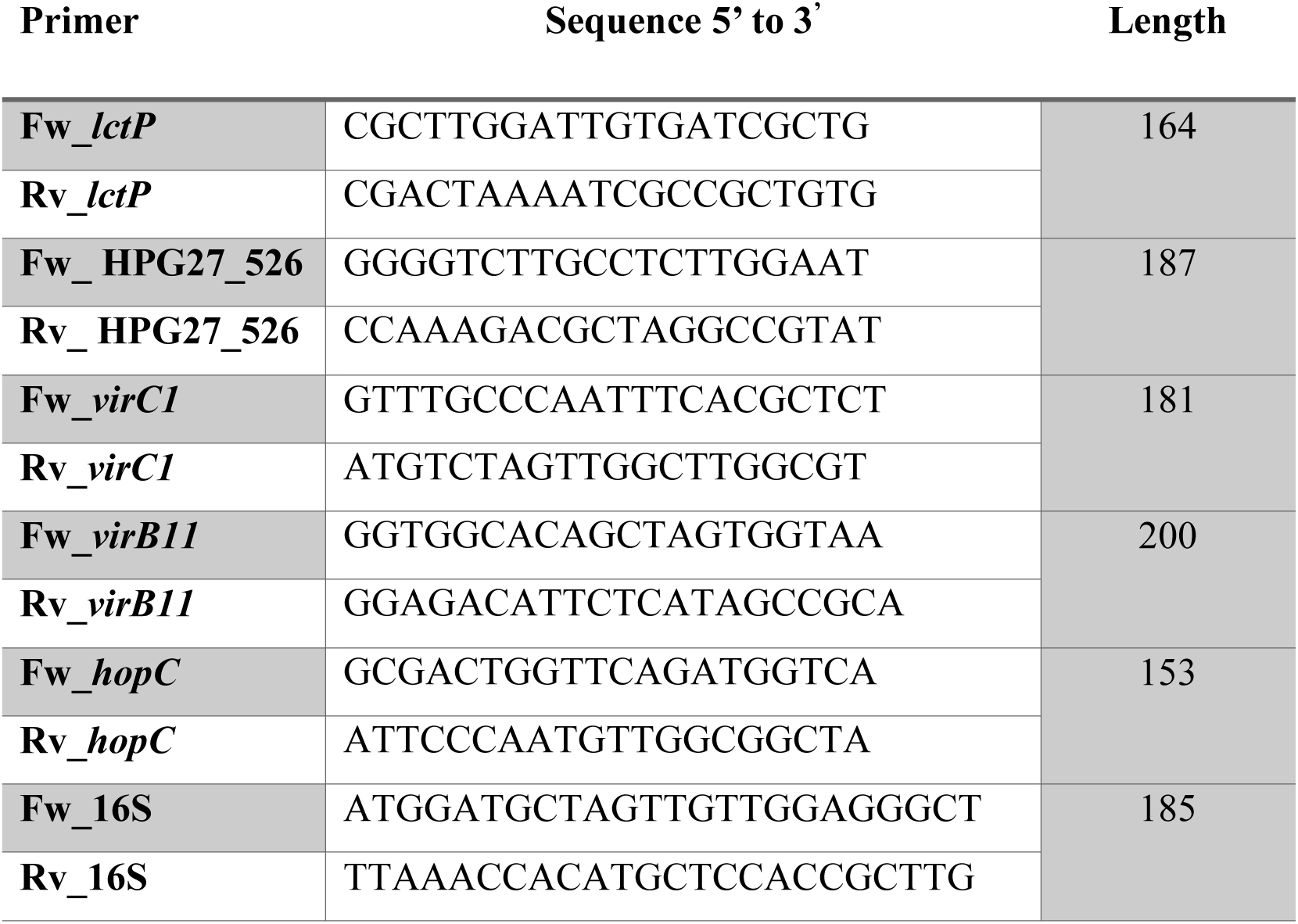
List of primers used in the present study for quantitative PCR.

**Supplemental_Table 2.**
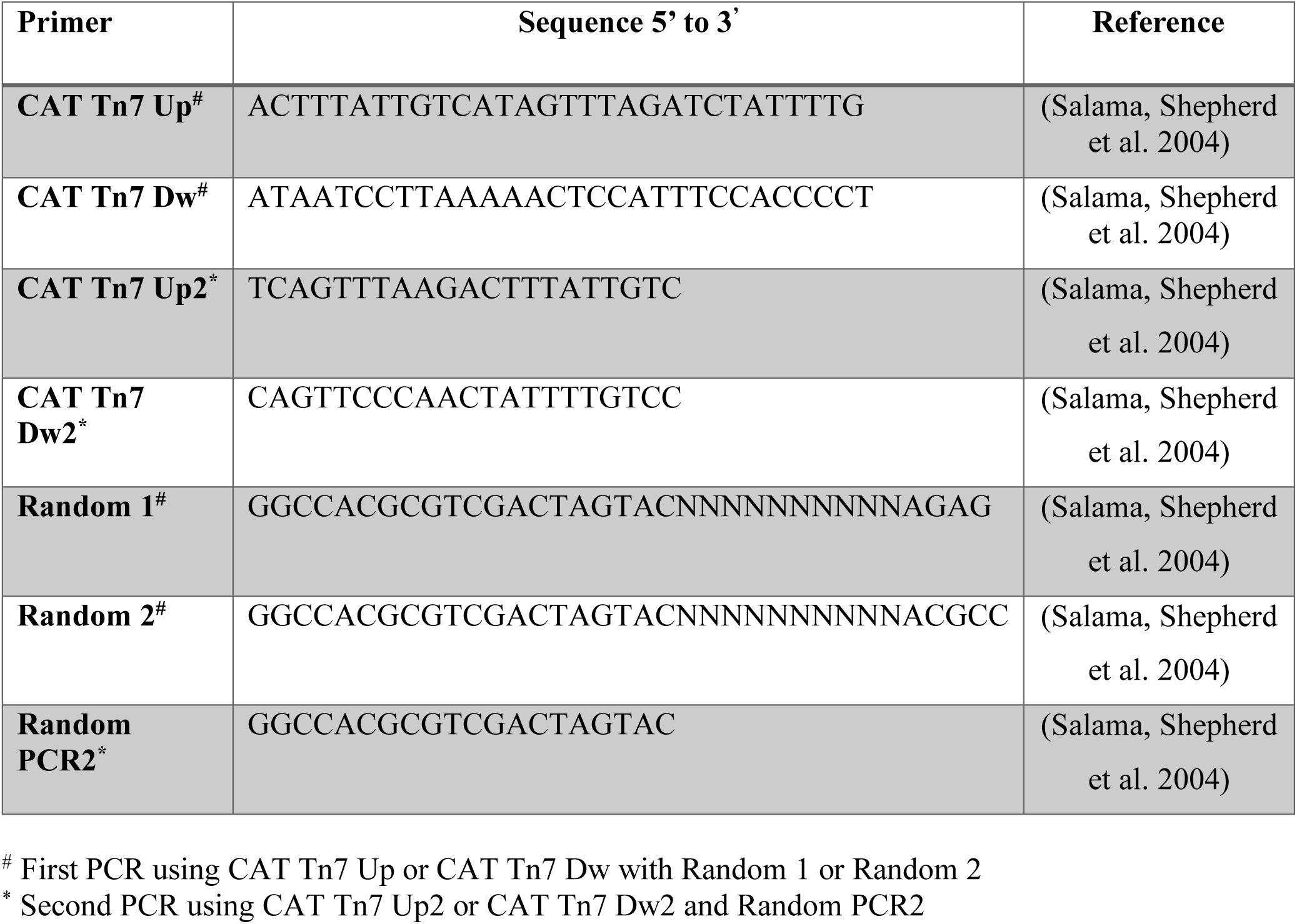
PCR Primers.

